# Flavoaffinins, elusive cellulose-binding natural products from an anaerobic bacterium

**DOI:** 10.1101/2025.02.08.637243

**Authors:** Duncan J. Kountz, Ruocheng Yu, Jessie H. Lee, Katherine N. Maloney, Emily P. Balskus

**Author notes:** Corresponding Author Emily P. Balskus Department of Chemistry and Chemical Biology, Harvard University, Cambridge, Massachusetts 02138, United States.

## Abstract

Cellulose is the most abundant polymer on earth. Many anaerobic cellulose-degrading bacteria produce uncharacterized yellow-orange, cellulose-binding pigments known as yellow affinity substances (here referred to as flavoaffinins) that are associated with cellulose degradation. Here, we isolate and structurally characterize the flavoaffinins from *Clostridium* (*Hungateiclostridium*) *thermocellum*, a key workhorse for the industrial conversion of cellulosic feedstocks to ethanol. Flavoaffinins represent an unprecedented structural juxtaposition of an aryl polyene chain with a hydroxy-diene γ-lactone. We also shed light on their biosynthesis using stable-isotope feeding experiments. This effort lays the groundwork for understanding the biological function(s) of the flavoaffinins and expands the limited number of natural products isolated from obligately anaerobic microbes.

---

Cellulose, the primary constituent of most plant cell walls^1^, is the planet’s most abundant polymer (1.5 × 10^12^ tons of biomass produced annually)^2^. As such, it is a major reservoir of fixed carbon, a potential renewable resource^3^, and the dominant microbial carbon source in most terrestrial ecosystems^4^. Anaerobic cellulose biodegradation to methane impacts global climate change^4^, while anaerobic cellulose degradation to short-chain fatty acids in ruminant livestock indirectly sustains humanity’s food supply^5^.

The genomes of anaerobic cellulose-degrading bacteria (cellulose fermenters) are enriched in natural product biosynthetic gene clusters (BGCs)^6^. However, few natural products have been identified from anaerobic bacteria^7-11^. In 1953, McBee^12^ isolated a thermophilic cellulose fermenter (*Clostridium thermocellum*, currently classified as *H. thermocellum*^13^) that produces a water-insoluble, yellow-orange pigment when grown on cellulose^12, 14^. In 1983, Ljungdahl *et al*. noticed that this pigment binds to both cellulose and endoglucanase, and proposed that it mediates attachment of the organism to cellulose fibrils^14^. This observation led them to name the pigment “yellow affinity substance.” Ljungdahl et al. also attempted to purify yellow affinity substance^14–16^; however, all isolation attempts have failed, likely due to the pigment’s oxygen and light sensitivity.

Here, we isolate and structurally characterize the flavoaffinins, a family of structurally unique cellulose-binding polyketides from *Clostridium thermocellum* that corresponds to the elusive yellow affinity substance. We also gain biosynthetic information from stable-isotopic feeding experiments, enabling future studies of the biosynthesis and biological functions of these unusual natural products.

To gain initial insights into the yellow affinity substance, we grew *C. thermocellum* DSM 1313 on cellulose, extracted the residual cellulose with acetone, and analyzed the extract using high-resolution tandem mass spectrometry (HR-MS/MS) with an in-line diode array UV-visible detector. This analysis revealed four major components, each with a λ_*max*_ of about 440 nm resembling the “carotenoid-like” UV-visible absorption band described previously (Figures 1 and S1).^14^ We observed these metabolites at *m/z* [M+H]^+^ 424.15, 450.17, 408.16, and 434.18, consistent with molecular formulae C_27_H_21_NO_4_, C_29_H_23_NO_4_, C_27_H_21_NO_3_, and C_29_H_23_NO_3_ (Table S1 and Figures S2 and S3). MS^2^ analysis revealed that each analyte produces prominent fragment ions with *m/z* 240.07 (C_14_H_10_NO_3_ ^+^) and 158.06 (C_10_H_8_NO^+^) (Figure S5). We named these metabolites flavoaffinins 423, 449, 407, and 433.

**Figure 1.**
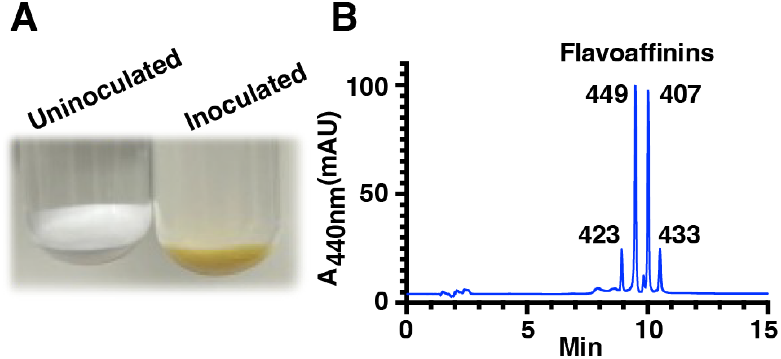
Analysis of *C. thermocellum* “yellow affinity substance” reveals four major components (flavoaffinins). (A) Cellulose sediments from uninoculated (left) and inoculated (right) growth medium following incubation at 60 °C for three days. (B) HPLC analysis of acetone extracts of cellulose sediment from a *C. thermocellum* culture grown in the dark. Peaks corresponding to flavoaffinins 423, 449, 407, and 433 are indicated.

We next optimized culturing and purification conditions to obtain sufficient quantities of flavoaffinins for characterization (Figure S4). As previously reported, we found that flavoaffinin production is induced by cellulose^14–16^. However, supplementation with 5 mM sodium acetate also induced flavoaffinin production during growth on cellobiose. Use of cellobiose permitted extraction of the flavoaffinins from the cell pellet with acetone. We used the flavoaffinins’ strong affinity for binding a microcrystalline cellulose affinity column later in the purification. We confirmed that flavoaffinins readily degrade under both light and oxygen (Figure S6); their stability in ambient light under anoxic conditions was not tested but should be considered when culturing *C. thermocellum*. Thus, we carried out the purification under red light and/or in an anaerobic chamber and obtained 3.9 mg of flavoaffinin 407 and 4.0 mg of flavoaffinin 449 from 324 L of *C. thermocellum* culture for structure determination.

Flavoaffinin 407 (**1**) (Figure 2, Table S2) was observed in HR-ESI-MS at *m/z* 408.1600, consistent with a molecular formula of C_27_H_21_NO_3_ (calcd 408.1605) and 18 degrees of unsaturation. The ^13^C NMR spectrum included 17 methine and 8 nonprotonated carbon signals at chemical shifts ranging from δ_c_ 97.9 to 170.9. COSY/TOCSY revealed four spin systems, including an aromatic methine proton at δ_H_ 7.95 (H-2) coupled to an exchangeable proton at δ_H_ 10.70 (H-1), a disubstituted benzene ring (H-5 through H-8), a phenyl group (H-24 through H-26), and a conjugated polyene chain (H-15 through H-22). HMBC correlations from H-2 to C-3, C-4 and C-9; from H-5 and H-7 to C-9; and from H-6 and H-8 to C-4 permitted assembly of the first two spin systems into an indole ring system substituted at the 3-position. At the other end of the molecule, HMBC correlations from H-21 to C-23, from H-22 to C-24, from H-24 to C-22 and from H-25 to C-23 connected the phenyl group to the aryl polyene chain. The dearth of proton-bearing carbons made elucidation of the remainder of the molecule challenging. However, key HMBC correlations from H-10 to C-4, C-11 and C-12 and from H-15 to C-12, C-13 and C-14 led us to propose the hydroxy-diene-γ-lactone moiety shown. The all-trans polyene configuration was determined based on the 3-bond proton coupling constants (*J*_*15,16*_ = 15.4 Hz; *J*_*16,17*_ = 11.5 Hz; *J*_*17,18*_ = 14.6 Hz; *J*_*18,19*_ = 11.4 Hz; *J*_*19,20*_ = 15.0 Hz; *J*_*21,22*_ = 15.6 Hz) and supported by NOESY correlations between H-15 and H-17; H-16 and H-18; H-19 and H-21; and H-21 and H-24. A NOESY correlation between H-5 and H-10 suggested a *s*-cis conformation for the C-3/C-10 bond. We assume the *Z-*configuration for the C-10/C-11 double bond for steric reasons. The conformation of the C-14/C-15 bond is not resolved by our data (Table S2).

**Figure 2.**
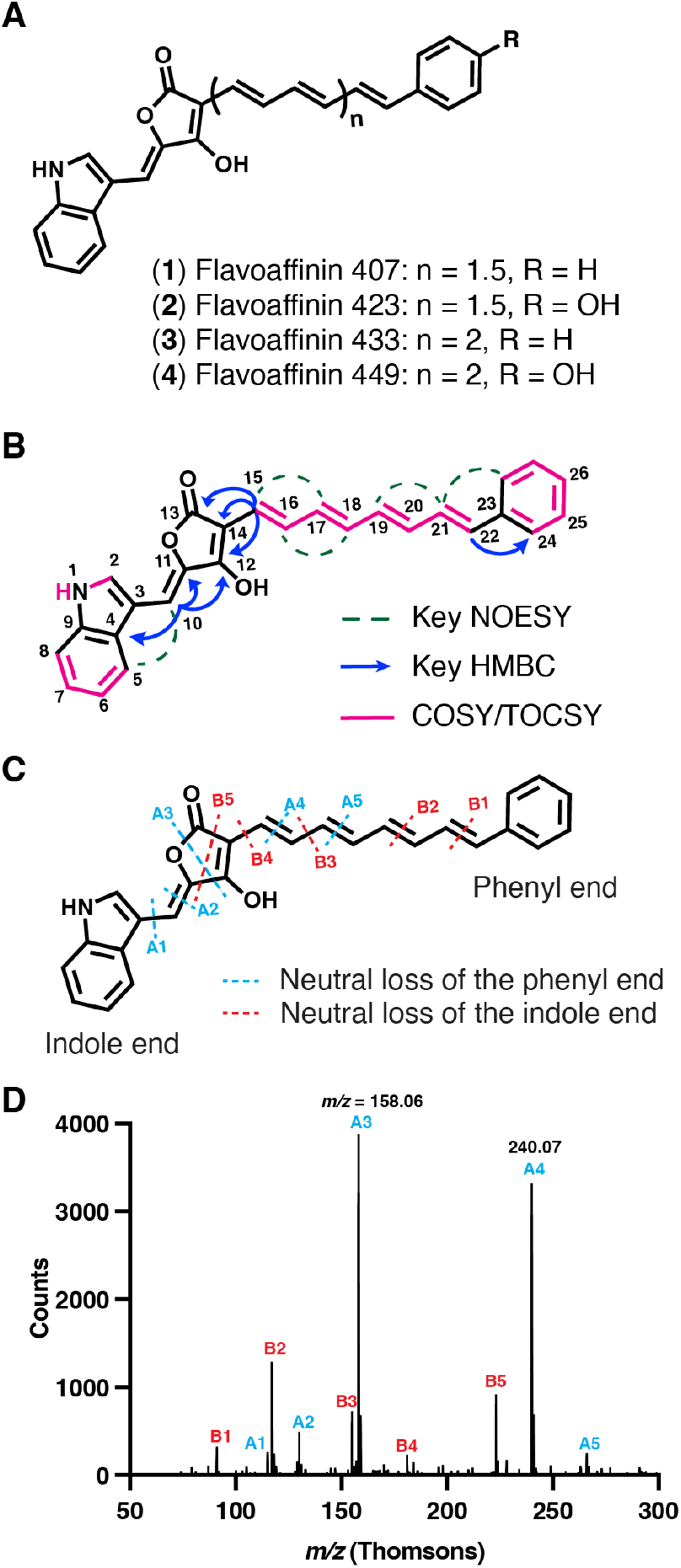
Structure elucidation of flavoaffinin 407 by NMR and HR-LC-MS/MS. (A) Proposed structures of flavoaffinins 407, 423, 433, and 449. (B) Key NOESY, HMBC, and COSY/TOCSY correlations observed for flavoaffinin 407. (C) HR-CID-MS^2^ fragmentation diagram for flavoaffinin 407 ([M + H]^+^ precursor ion). Blue and red lines indicate formation of fragment ions with neutral loss of the portion of the precursor ion connected to the phenyl group or indole ring, respectively. (D) HR-CID-MS^2^ of flavoaffinin 407 ([M + H]^+^ precursor ion.) Each peak is labeled with the name of the corresponding fragment ion given in panel C.

NMR data for flavoaffinin 449 (**4**) (Table S3) were similar to those of **1**, leading us to propose the same general structure consisting of a ‘head group’ composed of an indole linked to a hydroxy-diene-γ-lactone moiety, connected via an aryl polyene chain to a ‘tail group’. Apparent degradation of flavoaffinin 449 during anoxic storage at –70 °C may be attributed to partial oxidation (by disproportionation) of the phenolic hydroxy at C-26 followed by tautomerization through the C-12 enol as suggested by impaired resolution (Figures S22 and S23) of the signals in the polyene region of the spectrum. Nonetheless, we could tentatively assign the tail group as a *para*-substituted benzene ring (See NMR data in Table S3) on the basis of doublets at δ_H_ 7.29 (H-24) and 6.78 (H-25). The HR-MS/MS data discussed below suggests that the substituent is a *para*-hydroxy group.

The NMR-based structural assignments for flavoaffinins 407 and 449 are corroborated by HR-MS/MS data, which further permitted us to infer the structures of flavoaffinins 423 (**2**) and 433 (**3**) (Figure S5). All four flavoaffinins produce a C_14_H_10_NO_3_^+^ fragment ion (A4, see Figure 2D) corresponding to the head group. The presence of a monosubstituted indole ring was consistent with the production of indolium (C_8_H_8_N^+^, A1) and quinolinium (C_9_H_8_N^+^, A2) ions by all congeners. A polyene chain extending from the head group was supported by inspection of low-abundance ions, which revealed successive extension of the head group ion by multiple acetylene (C_2_H_2_) units (Figure S5).

Flavoaffinins 423 and 449 each produced abundant “tail group” hydroxytropylium ions (C_7_H_7_O^+^) suggestive of a hydrocarbon-substituted phenol. In the flavoaffinin 407 and 433 MS^2^ spectra, the hydroxytropylium was replaced by unsubstituted tropylium ions (C_7_H_7_ ^+^), suggesting a hydrocarbon substituted benzene ring in these congeners. In total, the HR-MS/MS data support the proposed structures of flavoaffinins 407 and 449, and suggest likely structures of flavoaffinins 423 and 433.

The flavoaffinins are unusual bacterial natural products. Hydroxy-diene-γ-lactones are rare in bacteria but common in plant and fungal natural products, including the aspulvinone^17,18^ pigments produced by *Aspergillus* spp. In the aspulvinones, the flanking aryl groups are phenyl or phenyl derivatives (Figure S36). Moreover, the non-indole aryl group of the flavoaffinins is separated from the central lactone by a polyene chain, which is unprecedented. Aryl polyenes are well-known antioxidant and/or photo-protectant bacterial polyketides that are often covalently-incorporated into the cell wall^19-21^. Although putative aryl polyene BGCs are widespread^22^, the flavoaffinins are (to our knowledge) the first aryl polyenes isolated from an obligately anaerobic bacterium. The flavoaffinin structures also provide potential insights into their cellulose-binding activity. Indole and indole-derived dyes bind cellulose^23,24^. Moreover, many proteins bind cellulose via multiple critical tryptophan residues^25-27^. These observations suggest the flavoaffinin indole moiety may be important for cellulose binding.

We next investigated the biosynthesis of the flavoaffinins. To test the hypothesis that the indole ring was derived from L-tryptophan, we performed stable isotopologue feeding experiments in *C. thermocellum*. Feeding *C. thermocellum* U-^13^C-L-tryptophan resulted in incorporation of all eleven carbons of tryptophan into all flavoaffinin congeners (Figure 3A and Figure S37B). Next, we assessed the origin of the terminal aryl group. As *C. thermocellum* lacks orthologs of genes involved in benzoate or hydroxybenzoate biosynthesis, we hypothesized that the aryl group would derive from L-phenylalanine (flavoaffinins 407 and 433) or L-tyrosine (flavoaffinins 423 and 449). Feeding experiments demonstrated incorporation of ring-D_5_-L-phenylalanine into flavoaffinins 407 and 433 (Figure 3B) and ring-D_4_-L-tyrosine into flavoaffinins 423 and 449 (Figure S37C). Finally, we hypothesized that the polyene would derive from acetate via the action of polyketide synthase (PKS) machinery. Feeding U-^13^C-acetate resulted in incorporation of up to three and four detected ketide units into flavoaffinins 407 and 449, respectively (Figure 3C and Figure S37D).

**Figure 3.**
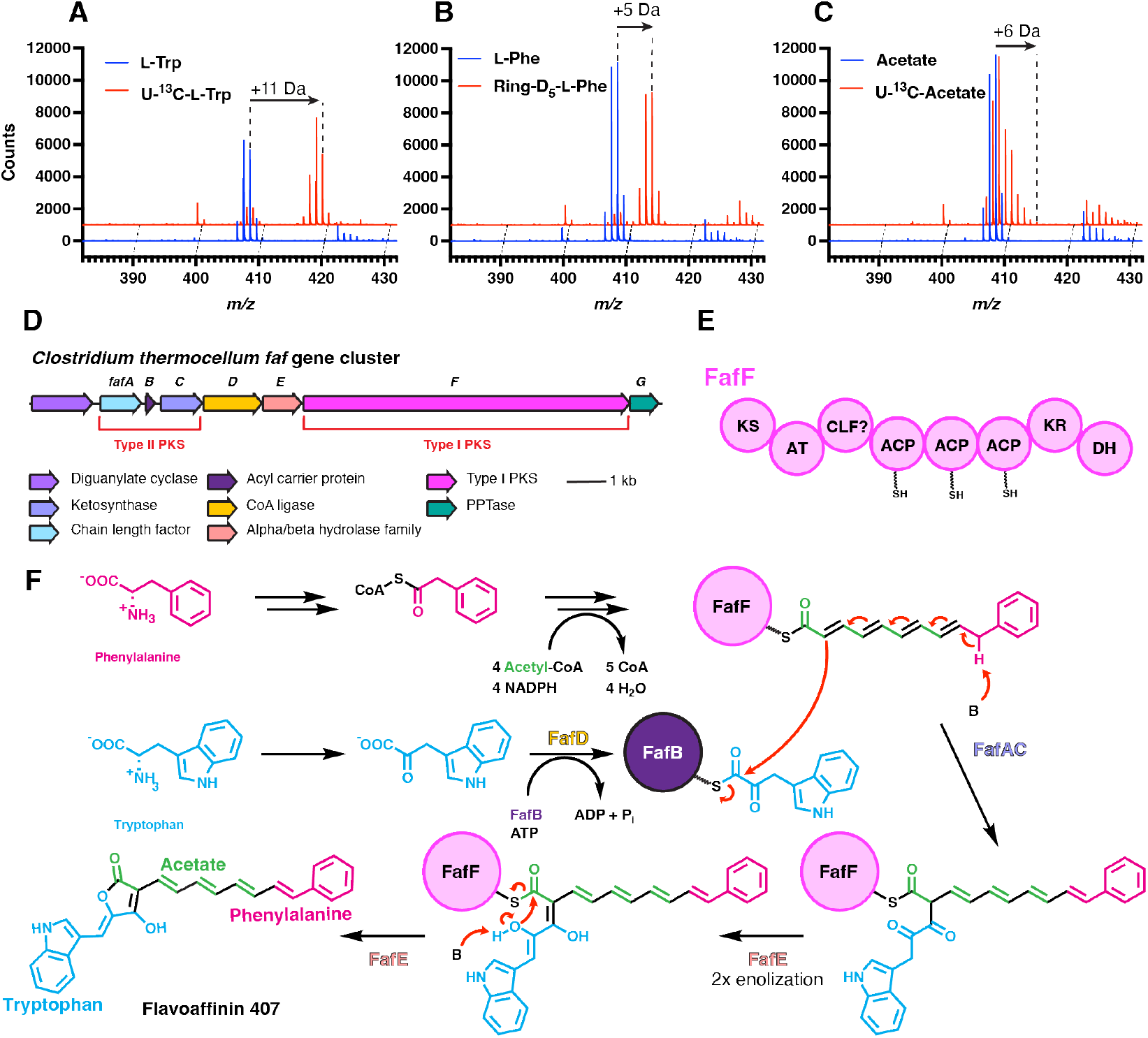
Flavoaffinins are biosynthesized by a polyketide synthase. (A) HR-LC-MS spectra of *C. thermocellum* fed either 0.25 mM unlabeled L-tryptophan (L-Trp) or 0.25 mM uniformly ^13^C-labeled L-tryptophan (U-^13^C-L-Trp). (B) HR-LC-MS spectra of *C. thermocellum* fed either 0.75 mM unlabeled L-phenylalanine (L-Phe) or 0.75 mM ring-perdeuterated L-phenylalanine (Ring-D5-L-Phe). HR-LC-MS spectra of *C. thermocellum* fed either 10 mM unlabeled sodium acetate (Acetate) or 10 mM uniformly ^13^C-labeled sodium acetate (U-^13^C-Acetate). For the ESI-positive mode mass spectra in (A)-(C) each flavoaffinin analyte was ionized by multiple pathways, including as a proton adduct, as a radical cation, and following in-source dehydrogenation, as previously described in ESI-MS of other polyenes^28^. (D) The putative flavoaffinin (*faf*) BGC (genes: Clo1313_2103-2096) from *C. thermocellum* DSM 1313 (E) The domain architecture of the type I PKS FafF. (F) Biosynthetic hypothesis for the flavoaffinins (also see Materials and Methods).. Abbreviations: PKS: polyketide synthase; PPTase: phosphopantetheinyl transferase; KS: ketosynthase; AT: acyltransferase; CLF: chain length factor; ACP: acyl carrier protein; KR: ketoreductase; DH: dehydratase.

These results allowed us to identify a putative flavoaffinin BGC. In particular, the labeling pattern observed when feeding acetate suggests the involvement of PKS machinery, as for all other aryl polyenes^20-22^. *C. thermocellum* DSM 1313 possesses a single PKS BGC (provisionally named *faf*) that is a strong candidate for flavoaffinin biosynthesis (Figure 3D). This BGC encodes a type I PKS (FafF) with a domain architecture consistent with aryl polyene biosynthesis (Figure 3E), alongside additional biosynthetic enzymes that could account for flavoaffinin production (Figure 3F). We suggest that the type I PKS (which lacks a reductase domain) synthesizes the thioester-tethered arylpolyene. In parallel, the CoA ligase homolog FafD catalyzes formation of a FafB-tethered indole-3-pyruvoyl group. These two thioesters could be condensed by the type II PKS FafAC to form a flavoaffinin. This proposal explains the incorporation of L-phenylalanine/L-tyrosine into the tail group, acetate into the polyene chain, and L-tryptophan into the head group.

To test the link between the *faf* gene cluster and flavoaffinin biosynthesis, we identified this gene cluster in additional bacterial genomes (see Materials and Methods) and examined two *faf*^*+*^ strains (*Pseudobacteroides cellulosolvens and Ruminiclostridium sufflavum*) for flavoaffinin production during growth on cellulose. Both strains produced flavoaffinin cogeners distinct from those produced by *C. thermocellum* (Figure S38). This finding strongly supports the involvement of the *faf* gene cluster in flavoaffinin production. While this manuscript was in review, Ishida et al. showed that disruption of the *fafF* gene eliminated flavoaffinin production in *C. thermocellum*^29^, verifying the function of this BGC. Only a handful of polyketides have been isolated from obligately anaerobic bacteria, making the flavoaffinins important additions to this class of natural products^7-10^.

Anaerobic cellulolytic bacteria are important industrial, ecological, agricultural, and environmental microbes. By elucidating the chemical structures of the enigmatic flavoaffinins, we now set the stage for investigating their biosynthesis and physiological functions, including potential impacts on cellulose degradation. Our findings also highlight anaerobic cellulolytic bacteria as an exciting source for the discovery of bioactive small molecules.

## ASSOCIATED CONTENT

### Supporting Information

Methods, NMR data for **1** and **4**, isotopic labeling of **4**, HR-MS and -MS^2^ data for **1**-**4** (PDF)

Raw NMR data files have been deposited on NP-MRD (https://np-mrd.org) and may be found under compound IDs NP0350641 (flavoaffinin 407) and NP0350642 (Flavoaffinin 449).

## AUTHOR INFORMATION

### Author Contributions

All authors contributed to the conceptualization, research, writing, and editing of the manuscript.

### Funding Sources

U.S. National Institutes of Health Training Grant #5T32GM095450

Howard Hughes Medical Institute (HHMI)-Gates Faculty Scholar Award (OPP1158186)

National Science foundation (NSF) Alan T. Waterman Award (CHE-20380529)

Howard Hughes Medical Institute

### Notes

The authors declare no competing financial interests.

## ACKNOWLEDGMENTS

D.J.K. acknowledges support from National Institutes of Health Training Grant #5T32GM095450. This work was supported by a Howard Hughes Medical Institute (HHMI)-Gates Faculty Scholar Award (OPP1158186) and a National Science foundation (NSF) Alan T. Waterman Award (CHE-20380529) to E.P.B. E.P.B. is Howard Hughes Medical Institute Investigator. We also note that this article is subject to HHMI’s Open Access to Publications policy. HHMI lab heads have previously granted a nonexclusive CC BY 4.0 license to the public and a sublicensable license to HHMI in their research articles. Pursuant to those licenses, the author-accepted manuscript of this article can be made freely available under a CC BY 4.0 license immediately upon publication. We thank Dr. Martin Daniel-Ivad and all other Balskus Laboratory members for comments and helpful discussion.

## ABBREVIATIONS

MS: mass spectrometry
LC, HPLC: high-performance liquid chromatography liquid chromatography
HR-MS/MS: high-resolution tandem mass spectrometry
UV: ultra-violet
NMR: nuclear magnetic resonance
COSY: ^1^H,^1^H-correlation spectroscopy
TOCSY: ^1^H,^1^H-total correlation spectroscopy
NOESY: nuclear Overhauser effect spectroscopy
HSQC: ^1^H,^13^C-heteronuclear single-quantum correlation spectroscopy
HMBC ^1^H,^13^: C-heteronuclear multiplebond correlation spectroscopy
BGC: biosynthetic gene cluster
PKS: polyketide synthase
ESI: electrospray ionization.

---

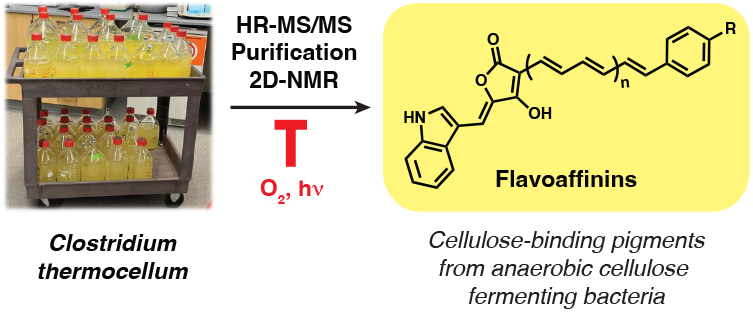

