## Supplemental Tables and Figures for "Flavoaffinins, elusive cellulose-binding natural products from an anaerobic bacterium"

**Supplemental Experimental Procedures**

**General Experimental Procedures.** All chemical and biological regents used in this study were from commercial sources (from Sigma-Aldrich, other than where noted), except for *C. thermocellum* DSM 1313 *Δhpt,* which was a kind gift of Dr. Adam Guss (Oak Ridge National Laboratory). Strain DSM 1313 *Δhpt* differs from the wildtype DSM 1313 strain by deletion of the *hpt* gene (locus tag CLO1313_RS14875), which encodes hypoxanthine phosphoribosyltransferase. This deletion was made previously to facilitate genetic manipulation and does not noticeably impact strain growth[1]. Because we do not expect this genetic modification to impact secondary metabolism, we suggest that *Δhpt* is appropriate for our study. Moreover, because this deletion is needed for genetic manipulation, its use here would simplify phenotypic comparisons made in future work that would involve genetic manipulation of *C. thermocellum*. Water from a MilliQ filter (IQ 7000) was used throughout the study.

**Cultivation of *C. thermocellum.*** *C. thermocellum* was cultivated in a modified formulation of the MTC medium[2], termed "mMTC" medium. The basal medium was composed of the following salts (per liter of basal medium): 1 mg resazurin sodium salt, 2.6 g sodium citrate monobasic, 1.5 g $NH_4Cl$, 2 g urea, 1 g $MgCl_2 \cdot 6 H_2O$, and 0.2 g $CaCl_2 \cdot 2 H_2O$. The basal media also contained 3 mL/L of a modified form of the Pfennig -Lippert trace-elements solution[3] that was in turn composed of the following components (per liter of trace-elements solution): 5 g ethylenediaminetetraacetic acid, 1.15 g $FeCl_2 \cdot 4 H_2O$, 47.4 mg $ZnCl_2$, 30 mg $MnCl_2 \cdot 4 H_2O$, 0.3 g boric acid, 0.2 g $CoCl_2 \cdot 6 H_2O$, 10 mg $CuCl_2 \cdot 2 H_2O$, 20 mg $NiCl_2 \cdot 6 H_2O$, 20 mg $Na_2MoO_4 \cdot 2 H_2O$. The pH of the trace elements solution was adjusted to 4.0 using HCl.

For small-scale (<120 mL media) cultures, basal medium was assembled and its pH adjusted to 7.2 with NaOH. The resulting medium was transferred to a vacuum flask, sealed with a rubber stopper and connected to an anaerobic gassing station where it was made anoxic by repeated cycles of exposure to vacuum with magnetic stirring, followed by flushes of 100% $N_2$. The flask was sealed, disconnected from the gassing station and transferred an anaerobic chamber (Coy Laboratories) with an atmosphere of 97.5% $N_2$/ 2.5 % $H_2$ and aliquoted into Balch tubes (10 mL of media/ ~28 mL volume tube) or 100 mL serum bottles (100 mL media/ ~120 mL bottle volume). The bottles or tubes were stoppered, crimped and had their atmospheres exchanged by brief cycles of vacuum followed by flushes of 100% $N_2$ using a gassing station. The vessels of media were autoclaved to a temperature of 121 °C for 20 minutes and cooled to room temperature. After autoclaving, the following sterile media components were added to the culture from anoxic solutions stored in serum bottles using anoxic, sterile tuberculin syringes (BD). To each 10 mL of media was added: 0.1 mL of 2.2 M potassium phosphate buffer (autoclaved under $N_2$), 0.05 mL 0.6 M L-cysteine • HCl (autoclaved under $N_2$), 0.05 mL 1 M $Na_2SO_4$, 0.05 mL 1 M sodium acetate (autoclaved under $N_2$), 0.2 mL of 0.5 M D-cellobiose (autoclaved under $N_2$), 0.2 mL of 10% (mass/volume) sodium bicarbonate (prepared under 20% $CO_2$/80% $N_2$) atmosphere and filter-sterilized using 0.2 micron Nylon syringe filters (Whatman) into a sterile serum bottle with a 100% $N_2$ atmosphere). Finally, immediately before inoculation, we added 0.05 mL filter-sterilized (0.2 micron Nylon syringe filters) vitamin solution/ 10 mL basal media. The vitamin solution (per liter) was composed of the following: 20 mg of biotin, 20 mg of folic acid, 30 mg of pyridoxine • HCl, 20 mg of thiamine • HCl, 10 mg of riboflavin, 20 mg of nicotinic acid, 20 mg of D-calcium-pantothenate, 10 mg of hydroxocobalamin • HCl, 10 mg of *p*-amino-benzoic acid, and 10 mg of lipoic acid. The pH of the vitamin solution was adjusted to pH 7.0 using NaOH, made anoxic under a 100% $N_2$ atmosphere and filter sterilized using 0.2-micron Nylon syringe filters into sterile serum bottles containing 100% $N_2$ atmospheres.

Large-scale cultures (>120 mL) were prepared as follows. The basal medium was assembled in the culture vessel and autoclaved to a temperature of 121 °C for 20–40 minutes. The cooled medium was supplemented per liter of medium with 20 mL of 0.5 M D-cellobiose, 5 mL of 1 M sodium sulfate, and 5 mL of 1 M sodium acetate (all sterilized by autoclaving). The medium was then made anoxic by bubbling with a sterile stream of $N_2$ at 20 PSI for 30 min/L media. The vessel was then sealed with a sterile septum and supplemented (per liter of media) with the following using a sterile, anoxic syringe: 2 mL 3 M L-cysteine • HCl, 20 mL 10% (mass/volume) sodium bicarbonate, and 5 mL vitamin solution (all prepared and sterilized as above).

*C. thermocellum* was cultivated in upright vessels in the dark at 60 °C with no shaking in an Innova 42R incubator (model r2R) for 24 hours. When cellulose was used as a carbon source, the basal media was autoclaved with 0.5% avicell PH-101 and no D-cellobiose was added after autoclaving. Cultures were exposed to ambient light only during brief daily examination for growth and (where applicable) during culture harvest. The flavoaffinins seemed to be stable to ambient light under culture conditions.

**Cultivation of *Pseudobacteroides cellulosolvens* and *Ruminiclostridium sufflavum*.** *P. cellulosolvens* and *R. sulffavum* were cultivated in the same manner as *C. thermocellum*, but with the following differences.

*P. cellulosolvens* DSM 2933 was grown in a modified form of "*Pseudobacteroides cellulosolvens* medium" (DSMZ medium 315). The basal medium contained the following (per L): 0.74 g $NH_4Cl$, 0.1 g $MgCl_2$ • 6 $H_2O$, 0.6 g $CaCl_2$ • 2 $H_2O$, 3 mL Pfennig-Lippert trace-elements solution[3] (for composition, see above *Cultivation of C. thermocellum*), and 1 mg resazurin sodium salt. The basal media was made anoxic and aliquoted as specified above (*Cultivation of C. thermocellum)* for mMTC media and then autoclaved. After the media had cooled, the following sterile media components were added to the culture from anoxic solutions stored in serum bottles using anoxic, sterile tuberculin syringes (BD). To each 10 mL of media was added: 0.1 mL of 2.2 M potassium phosphate buffer (autoclaved under $N_2$), 0.05 mL 0.6 M L-cysteine • HCl (autoclaved under $N_2$), 0.05 mL 0.2 M $Na_2S$ (autoclaved under $N_2$), 0.05 mL 1 M $Na_2SO_4$, 0.2 mL of 0.5 M D-cellobiose (autoclaved under $N_2$), 0.2 mL of 10% (mass/volume) sodium bicarbonate (prepared under 20% $CO_2$/80% $N_2$) atmosphere and filter-sterilized using 0.2 micron Nylon syringe filters (Whatman) into a sterile serum bottle with a 100% $N_2$ atmosphere). Finally, immediately before inoculation, we added 0.05 mL filter-sterilized (0.2 micron Nylon syringe filters) vitamin solution/ 10 mL basal media (see *Cultivation of C. thermocellum* above for composition of the vitamin solution). When cellulose was used as a carbon source, the basal media was autoclaved with 0.5% avicell PH-101 and no D-cellobiose was added after autoclaving. *P. cellulosolvens* was cultivated in upright vessels in the dark at 30 °C with no shaking in an Innova 42R incubator (model r2R). While growth was faster at 37 °C, pigment production was weak or null at this temperature.

*R. sufflavum* DSM 19573 was grown in a modified form of "PY + X medium" (DSMZ medium 104b). The basal medium contained (per L): 5 g tryptone, 5 g peptone, and 10 g yeast extract (all from Research Products International), 0.1 g $CaCl_2 \cdot 2 H_2O$, 0.02 g $MgSO_4 \cdot 7 H_2O$, and 2 g NaCl. The basal media was made anoxic and aliquoted as specified above (*Cultivation of C. thermocellum)* for mMTC media and then autoclaved. After the media had cooled, the following sterile media components were added to the culture from anoxic solutions stored in serum bottles using anoxic, sterile tuberculin syringes (BD). To each 10 mL of media was added: 0.1 mL of 2.2 M potassium phosphate buffer (autoclaved under $N_2$), 0.05 mL 0.6 M L-cysteine • HCl (autoclaved under $N_2$), 0.2 mL of 0.5 M D-cellobiose (autoclaved under $N_2$), 0.2 mL of 10% (mass/volume) sodium bicarbonate (prepared under 20% $CO_2$/80% $N_2$) atmosphere and filter-sterilized using 0.2 micron Nylon syringe filters (Whatman) into a sterile serum bottle with a 100% $N_2$ atmosphere). Finally, immediately before inoculation, we added 0.05 mL filter-sterilized (0.2 micron Nylon syringe filters) vitamin solution/ 10 mL basal media (see *Cultivation of C. thermocellum* above for composition of the vitamin solution). When cellulose was used as a carbon source, the basal media was autoclaved with 0.5% avicell PH-101 and no D-cellobiose was added after autoclaving. *P. cellulosolvens* was cultivated in upright vessels in the dark at 30 °C with no shaking in an Innova 42R incubator (model r2R).

**High-Resolution Liquid Chromatography-Mass Spectrometry (HR-LC-MS) analysis.** HR-ESI-MS and MS/MS analysis was conducted on an Agilent Technologies 6530 Accurate-Mass Q-TOF equipped with a dual Agilent Jet Stream Electrospray Ionization source. Chromatography was performed using a Hypersil Gold™ aQ column (Thermo Fisher Scientific; dimensions: 250 × 10 mm; particle size: 3 μm). All solvents were LC-MS grade (Sigma Aldrich) and were supplemented with 0.1% LC-MS grade formic acid (Sigma Aldrich). After a 2 minute wash of 1:1 acetonitrile:water supplemented with 0.1% formic acid, the chromatography method ran from 1:1 to 19:1 acetonitrile:water over 12 minutes. The gradient was followed by a 5 minute wash with 19:1 acetonitrile:water before reequilibration in 1:1 acetonitrile:water. UV-visible spectral profiles of analytes were collected following the chromatography step via a diode array cell (Agilent Technologies 1260 Infinity) in line with the mass spectrometer source. Positive mode ionization was used with a drying gas ($N_2$) temperature of 275 °C (flow rate 10 L/min). The nebulizer was set to 45 psi, and the sheath gas temperature and flow rate were 250 °C and 11 L/min, respectively. The Dual AJS ESI source voltage was 4000 V with a nozzle voltage of 500 V. MS TOF parameters for auto MS/MS were set as follows. Fragmentor: 125 V; Skimmer: 65 V; Oct 1 RD Vpp: 750 V. MS transients were scanned between $m/z$ = 100 and $m/z$ = 700. Acquisition rate and time were one spectrum/second and 1000 ms/spectrum, respectively, with 13637 transients/spectrum. For auto MS/MS, transients were scanned between $m/z$ = 30 and $m/z$ = 700 with the same acquisition rate and time as at the MS level, but with 13487 transients/spectrum. Isolation width was set to ~4 *m/z*. Collision energy was 20 units. Precursor selection parameters were as follows. Max precursor per cycle: 5; predictor threshold: 200 (Ref. threshold 0.01%). Active exclusion of precursor ions was enacted after 3 spectra and the exclusion was released after 0.5 minutes. Mass error tolerance for iterative MS/MS was ± 20 ppm and RT exclusion tolerance was 0.2 (-min) an 0.2 (+min). We used the “peptides” isotope model with a purity stringency of 100% and purity cutoff of 30%. The instrument was calibrated monthly using Agilent ESI-L LCMS Tuning solution (part number: G1969-85000), and tuned prior to each experimental run. During the runs, the time-of-flight tube was internally calibrated by infusion of purine (*m/z* = 121.05087) and HP-921 (hexakis(1H, 1H, 3H-tetrafluoropropoxy)phosphazine), *m/z* = 922.009798).

**Isolation of the flavoaffinins.** 324 L of *C. thermocellum* was grown as described above in 2 L and 5 L Duran GL 45 laboratory bottles sealed with anaerobic flange stoppers (Chemglass) and aperture screw caps (DWK Life Sciences). The 2 and 5 L bottles contained 1.5 and 4 L (respectively) of mMTC + cellobiose medium supplemented with 5 mM sodium acetate. Starter cultures of 100 mL were grown for 24 hours in mMTC + cellobiose medium supplemented with 5 mM sodium acetate and used to inoculate the large-scale cultures composed of the same medium with 3 mL of inoculum into 1 L of medium. The resulting cultures were incubated at 60 °C for 24 hours and harvested under an ambient atmosphere by centrifugation. Subsequent steps were always performed in the absence of direct or indirect sunlight, and often under red light (and otherwise in the dark.)

The combined cell pellet from all of the cultures was extracted with acetone (Sigma Aldrich, ACS grade) and the suspension was centrifuged to pellet cell debris. The extraction of the cell pellet was repeated until little or no color remained in the supernatant. This provided ~3 L of acetone extract. The acetone extract was concentrated using a rotary evaporator to provide a mixture of solids and oil that was redissolved in 1 L of 1:5 reagent grade ethanol:water. The resulting solution was loaded onto a series of three 10 g Sep-Pak (Waters) C18 solid-phase extraction cartridges, which were then washed with 5-10 volumes of 1:5 reagent grade ethanol:water. The flavoaffinins were then eluted with 100% HPLC-grade acetone. This acetone solution was concentrated to an oil using a rotary evaporator. 100 g of microcrystalline cellulose suspended in 500 mL of HPLC-grade ethanol were then added to the flask containing the oil and dried using a rotary evaporator until only the damp cellulose remained. 300 mL of water were then added and the evaporation was repeated. The resulting cellulose was resuspended in ~1 L of water and poured into a column. The column was washed with 1 L of 100% HPLC-grade methanol, then the flavoaffinins were eluted with 1 L of HPLC-grade acetone containing 1 g of sodium hydroxide. The eluate was concentrated about 4-fold on a rotary evaporator, then mixed with 1 L of water and concentrated until about 500 mL remained. This solution was extracted twice with 1 L of portions of HPLC-grade ethyl acetate. The organic layers were collected and dried on a rotary evaporator.

The residue was resuspended in 2 mL of LC-MS grade methanol and injected (four runs, each with a 0.5 mL injection volume) onto a semi-preparative Hypersil Gold™ aQ column (Thermo-Fisher Scientific; dimensions: 250 × 10 mm; particle size: 5 μ) using a

Dionex 3000 preparatory HPLC system (Thermo-Fisher Scientific). The chromatographic method was composed of a 20 minute isocratic flow with 1:1 water:acetonitrile, followed by a 100 minute linear gradient from 1:1 water:acetonitrile to 1:99 water:acetonitrile. 0.1% formic acid was included in both solvents and the flow rate was 4 mL/minute. The peaks corresponding to flavoaffinins 407 and 449 were collected and the identities of the analytes confirmed by high-resolution LC-MS on a Q-TOF as described above. The fractions containing each congener were pooled, dried using a rotary evaporator, and then on a Centrivap (EZ-2.3 Elite Evaporation System) at 40 °C in preparation for NMR. This procedure yielded 3.9 and 4.0 mg of flavoaffinins 407 and 449, respectively from 324 L of culture.

**NMR-based structural elucidation of the flavoaffinins.** The dried flavoaffinin 407 and 449 preparations (see above) were transferred to an Mbraun anaerobic chamber that was maintained under an atmosphere of 100% $N_2$. Inside the chamber each sample was dissolved in 500 μL of $(CD_3)_2CO$ (Cambridge Isotopes) stored in a sealed ampule under argon. Each solution was transferred to a 5 mm Low Pressure/Vacuum (LPV) 600 MHz NMR tube which was then sealed with its Teflon plug and removed from the chamber.

NMR spectra were recorded on a Bruker Avance II 600 MHz Bio-molecular NMR system housed at the Harvard Medical School Biomolecular NMR Facility and DFCI NMR Core. Chemical shifts are reported in ppm relative to the $(CD_3)_2CO$ solvent residuals ($\delta$ = 29.84 and 2.05 ppm for $^{13}C$ and $^{1}H$ spectra, respectively). A 400 MHz spectrum of flavoaffinin 449 was acquired on a JEOL ECZ400S NMR spectrometer.

A contaminant (2,2'-methylenebis(6-*tert*-butyl-*p*-cresol), (**5**) was present in both flavoaffinin NMR samples. The identity of this contaminant was confirmed by determining the $^{1}H$ NMR spectrum of an authentic standard (purchased from TCI Chemicals) using a Bruker AVANCE NEO 400B NMR spectrometer (Table S4).

**Flavoaffinin 407 (1)**: orange-red amorphous solid; UV (1:1 water:ACN + 0.1% formic acid) $\lambda_{max}$ 362, 442 nm (Figure S1); $^{1}H$, $^{13}C$ and 2D NMR spectroscopic data, see Table S1; (+)-HRESIMS *m/z* 408.1600 [M + H]$^+$ (calculated for $C_{27}H_{22}NO_3^+$, 408.1594)

**Flavoaffinin 449 (4)**: orange-red amorphous solid; UV (1:1 water:ACN + 0.1% formic acid) $\lambda_{max}$ 378, 452 nm (Figure S1); $^{1}H$, $^{13}C$ and 2D NMR spectroscopic data, see Table S2; (+)-HRESIMS *m/z* 450.1712 [M + H]$^+$ (calculated for $C_{29}H_{24}NO_4^+$, 450.1700)

**Feeding experiments with isotopically-labeled substrates.** Balch tubes containing 10 mL of mMTC-cellobiose medium prepared as described above (but lacking sodium acetate) were amended with anoxic solutions of isotopically-labeled or unlabeled substrates noted below, sterilized with 0.2-micron polyethanesulfone syringe filters (VWR International). For each labeling experiment, the test group consisted of three cultures, each supplemented with 0.1 mL of 25 mM U-$^{13}C$-L-tryptophan, while the control group was composed of three cultures, each supplemented with 0.1 mL of 25 mM unlabeled L-tryptophan. Analogous experiments were performed with addition of 0.3 mL of 25 mM unlabeled or ring-$D_4$-L-tyrosine, 0.3 mL of 25 mM unlabeled or ring-$D_4$-L-phenylalanine, or 0.2 mL of 0.5 M unlabeled or U-$^{13}C$-labeled sodium acetate. Unlabeled compounds were obtained from Sigma-Aldrich, while labeled compounds were obtained from Cambridge Isotope Laboratories. The cultures were inoculated with 0.1 mL of a 24-hour grown starter culture and incubated at 60 °C for 24 hours. The cultures were then harvested by centrifugation, and the resulting pellets washed with 1 mL of 44 mM potassium phosphate, pH 7.2 by resuspension and re-centrifugation. The pellets were extracted with 300 μL of LC-MS grade acetonitrile and centrifuged for 10 min at 16,100 × g to pellet cell debris. The supernatants were transferred to new tubes and recentrifuged. The final supernatants were analyzed on a Q-TOF as described above.

**Bioinformatic identification of a putative flavoaffinin (*faf*) biosynthetic gene cluster.** The published *C. thermocellum* DSM 1313 genome sequence (RefSeq accession: NC_017304) was queried using antiSMASH 7.0[4]. Domain assignments were made using antiSMASH 7.0 and manually using InterPro. Additional *faf* gene clusters were identified by searching the NCBI ClusteredNR database using BLASTP with FafD (WP_003512157.1/Clo1313_2099 gene product) as a query. The genomic contexts of hits with E values below E-80 were inspected manually to confirm the presence gene encoding a FafF ortholog in the same gene cluster cluster as the encoded FafD ortholog. The organisms encoding both a FafD ortholog and a FafF ortholog in the same gene cluster were then selected and their genomes downloaded. The predicted proteomes were used to generate a custom protein database that was searched using BLASTP with each gene product (FafA-G) in the *C. thermocellum faf* gene cluster. It was found that each of the nine organisms organisms encoded complete or nearly complete *faf* gene clusters. In two cases (*A. clariflavus* and *A. cellulolyticus*) it was found that FafE was encoded in a separate gene cluster.

**Biosynthetic hypothesis for the flavoaffinins.** Phenylacetyl-CoA can be readily produced by oxidative decarboxylation of phenylalanine via phenylpyruvate. The oxidative decarboxylation is likely catalyzed by the arylpyruvate:ferredoxin oxidoreductase IorAB (genes: Clo1313_1616-1615). The oxidative deamination of L-phenylalanine to phenylpyruvate is likely catalyzed by an α-aminoacid aminotransferase encoded by the Clo1313_1619 gene nearby *iorAB*. The Clo1313_1619 gene product could also be responsible for deaminating L-tryptophan to indole-3-pyruvate. Flavoaffinins containing a *p*-hydroxyphenyl group as a part of the arylpolyene moiety incorporate this ring for tyrosine as shown in Figure S37. The variable polyene chain length observed across flavoaffinin congeners is likely due to the iterative action of polyene chain elongation by FafF.

**Table S1.** Summary of HR-ESI-MS measurements of the flavoaffinin *m/z* values

| Analyte | Measured *m/z* | Formula (Neutral molecule) | *m/z* error (ppm) |
|---|---|---|---|
| **Flavoaffinin 423** | 424.1532 | $C_{27}H_{21}NO_4$ ($[M + H]^+$ *m/z* = 424.1543) | - 2.67 |
| **Flavoaffinin 449** | 450.1712 | $C_{29}H_{23}NO_4$ ($[M + H]^+$ *m/z* = 450.1700) | 2.70 |
| **Flavoaffinin 407** | 408.1600 | $C_{27}H_{21}NO_3$ ($[M + H]^+$ *m/z* = 408.1605) | 1.42 |
| **Flavoaffinin 433** | 434.1748 | $C_{29}H_{23}NO_3$ ($[M + H]^+$ *m/z* = 434.1751) | - 0.62 |

**Table S2.** Table of NMR spectroscopic data for flavoaffinin 407 in $(CD_3)_2CO$

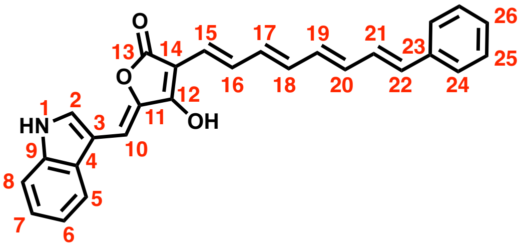

**Flavoaffinin 407 (1)**

| Position | ${}^{a}\delta_C$, type | ${}^{b}\delta_H$, mult. (*J* in Hz) | COSY | TOCSY | NOESY | HMBC (J = 8 Hz) |
|---|---|---|---|---|---|---|
| 1 | - | 10.7 (s, 1H) | 2 | - | - | - |
| 2 | 128.4, CH | 7.95 (d, 2.0, 1H) | 1 | - | - | 3, 4, 9 |
| 3 | 111.0, C | - | - | - | - | - |
| 4 | 128.0, C | - | - | - | - | - |
| 5 | 119.4, CH | 7.91 (d, 7.8, 1H) | 6 | 6, 7, 8 | 6, 10 | 7, 9 |
| 6 | 120.9, CH | 7.10 (ddd, 7.9, 7.0, 1.0, 1H) | 5 | 5, 7, 8 | 5 | 4, 8 |
| 7 | 122.9, CH | 7.16 (ddd, 8.1, 7.0, 1.2, 1H) | 8 | 5, 6, 8 | 8 | 5, 9 |
| 8 | 112.5, CH | 7.47 (d, 8.1, 1H) | 7 | 5, 6, 7 | 7 | 4, 6 |
| 9 | 137.1, C | - | - | - | - | - |
| 10 | 98.3, CH | 7.00 (s, 1H) | - | - | 5 | 4, 11, 12 |
| 11 | 144.0, C | - | - | - | - | - |
| 12 | 170.9, C | - | - | - | - | - |
| 13 | 170.2, C | - | - | - | - | - |
| 14 | 97.9, C | - | - | - | - | - |
| 15 | 124.1, CH | 6.71 (d, 15.4, 1H) | 16 | 16, 17, 18 | 17 | 16, 17, 12, 13 |
| 16 | 127.0, CH | 7.27 (m, 1H) | 15, 17 | 15, 17, 18, 19 | 18 | 14 |
| 17 | 137.6, CH | 6.49 (dd, 14.6, 11.5, 1H) | 16 | 15, 16 | 15 | 15 |
| 18 | 130.7, CH | 6.35 (dd, 14.6, 10.6, 1H) | c | c | 16 | 17, 19 |
| 19 | 135.8, CH | 6.48 (dd, 15.0, 11.4, 1H) | c | c | 21 | 21 |
| 20 | 132.1, CH | 6.40 (dd, 15.0, 10.5, 1H) | c | c | c | 22 |
| 21 | 130.0, CH | 6.95 (dd, 15.6, 10.5, 1H) | c | c | 19, 24 | 19, 23 |
| 22 | 131.8, CH | 6.55 (d, 15.6, 1H) | c | c | 24 | 20, 24 |
| 23 | 138.8, C | - | - | - | - | - |
| 24 | 127.0, CH | 7.43 (d, 7.9, 2H) | 25 | 25, 26 | 21, 22, 25 | 22, 26 |
| 25 | 129.4, CH | 7.28 (t, 7.1, 2H) | 24, 26 | 24, 26 | 24 | 23, 24 |
| 26 | 127.9, CH | 7.17 (tt, 6.7, 1.2, 1H) | 25 | 24, 25 | c | 24 |

[a]Recorded at 125 MHz; referenced to residual $({}^{13}CD_3)_2CO$ at 29.8406 ppm
[b]Recorded at 600 MHz; referenced to residual $(CHD_2)CD_3CO$ at 2.0500 ppm
[c]Overlapping

**Table S3.** Table of NMR spectroscopic data for flavoaffinin 449 in $(CD_3)_2CO$. Due to extensively overlapping and broadened signals, we could not resolve the resonances corresponding to most of the polyene chain.

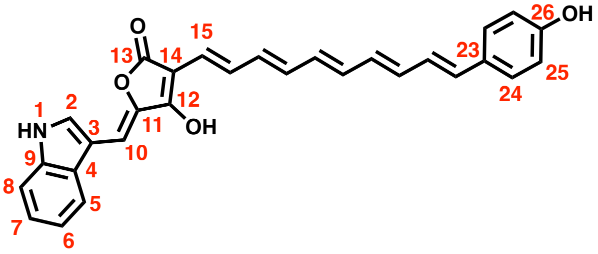

**Flavoaffinin 449 (4)**

| Position | $^a\delta_C$, type | $^b\delta_H$, mult. (*J* in Hz) | COSY | TOCSY | HMBC (J = 8 Hz) |
|---|---|---|---|---|---|
| 1 | - | 10.63 (s, 1H) | 2 | - | - |
| 2 | 127.0, CH | 7.92 (d, 2.1, 1H) | 1 | - | 3, 4, 9 |
| 3 | 111.5, C | - | - | - | - |
| 4 | 128.0, C | - | - | - | - |
| 5 | 118.5, CH | 7.88 (d, 7.7, 1H) | 6 | 6, 7, 8 | 3, 4, 7, 9 |
| 6 | 119.8, CH | 7.09 (ddd, 8.2, 7.0, 0.8, 1H) | 5, 7 | 5, 7, 8 | 4, 8 |
| 7 | 121.9, CH | 7.15 (ddd, 8.1, 7.2, 0.8, 1H) | 6, 8 | 5, 6, 8 | 6, 9 |
| 8 | 112.2, CH | 7.45 (d, 8.0, 1H) | 7 | 5, 6, 7 | 4, 6 |
| 9 | 137.0, C | - | - | - | - |
| 10 | 96.2, CH | 6.9 (s, 1H) | - | - | 4, 11 |
| 11 | 145.1, C | - | - | - | - |
| 12 | c | - | - | - | - |
| 13 | c | - | - | - | - |
| 14 | c | - | - | - | - |
| 15 | 124.9, CH | 6.66 (d, 15.8 1H) | d | d | - |
| 23 | 116.0, C | - | - | - | - |
| 24 | 127.7, CH | 7.29 (broad, 2H) | 25 | 25 | - |
| 25 | 115.5, CH | 6.78 (d, 8.2, 2H) | 24 | 24 | 23, 26 |
| 26 | 158.1, C | - | - | - | - |

[a]Recorded at 125 MHz; referenced to residual $(^{13}CD_3)_2CO$ at 29.8406 ppm
[b]Recorded at 600 MHz; referenced to residual $(CHD_2)CD_3CO$ at 2.0500 ppm
[c]not observed
[d]correlates with a poorly-resolved resonance at $\delta_H$ = 7.21 ppm

**Table S4:** Table of NMR spectroscopic data for an authentic standard of 2,2'-methylenebis(6-tert-butyl-p-cresol) (**5**), a contaminant in the flavoaffinin 407 and 449 NMR samples, in $(CD_3)_2CO$

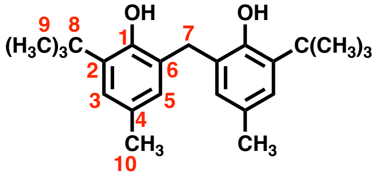

**Contaminant (5)**

| Position | Type | $^a\delta_H$, mult. (*J* in Hz) |
|---|---|---|
| 1 | OH | 7.55 (br s, 2H) |
| 2 | C | - |
| 3 | CH | 6.94 (d, 2.2, 2H) |
| 4 | C | - |
| 5 | CH | 6.84 (dd, 2.2, 0.9, 2H) |
| 6 | C | - |
| 7 | $CH_2$ | 3.90 (s, 2H) |
| 8 | C | - |
| 9 | $CH_3$ | 1.38 (s,18H) |
| 10 | $CH_3$ | 2.18 (s, 6H) |

[a]Recorded at 400 MHz; referenced to residual $(CHD_2)CD_3CO$ at 2.0500 ppm

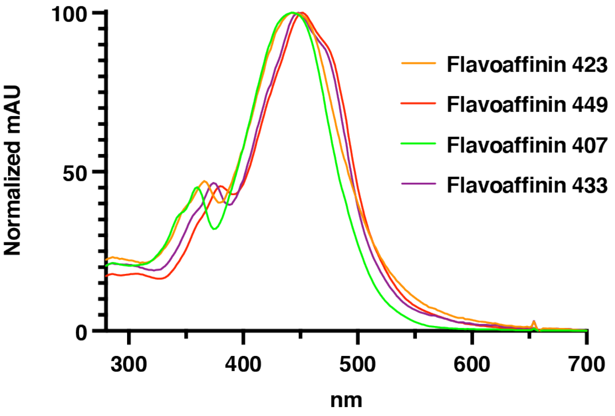

**Figure S1.** Ultraviolet-Visible (UV-Vis) spectra of the flavoaffinins. Spectra were collected following elution from an HPLC system with an in-line diode array detector. The solvent mixture at the time of each elution was about 75% acetonitrile, 25% water, and 0.1% formic acid. For comparison, the spectra were normalized so that the $\lambda_{max}$ of the 400-500 nm absorption band was 100 mAU.

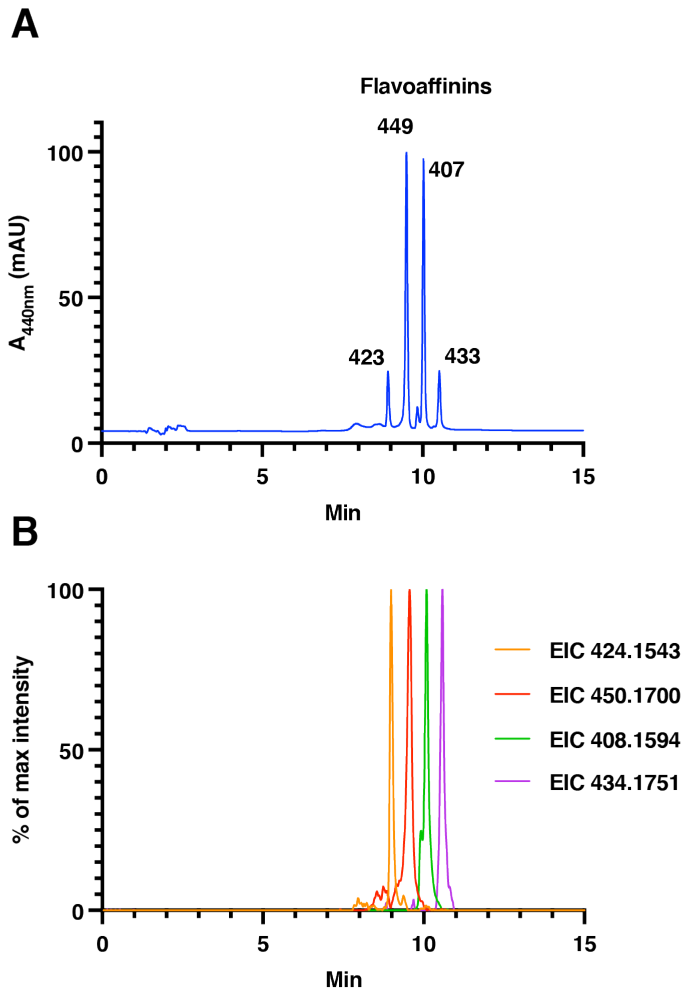

**Figure S2.** Correlation of the retention times of the flavoaffinin peaks in the $A_{440nm}$ trace (A) with the extracted ion chromatogram (EIC) peaks for the inferred *m/z* value for each congener (B). Note that Panel A is reproduced from Figure 1B in the main text. In B, the EICs are scaled to have the same maximum intensity for convenient comparison.

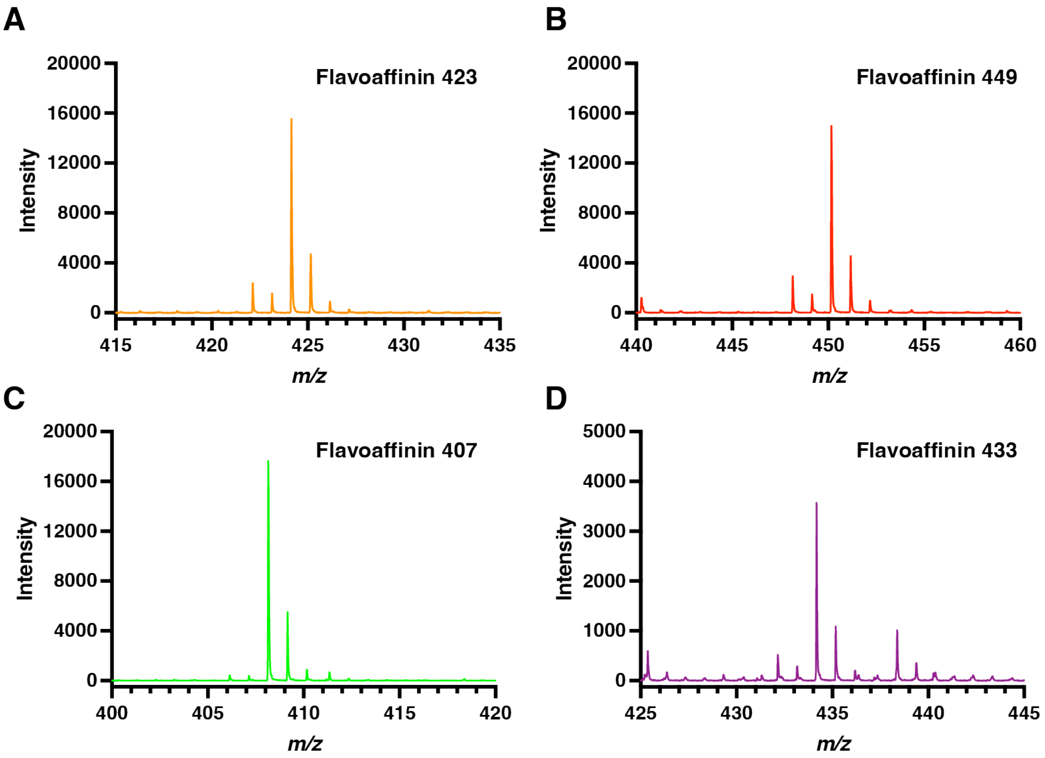

**Figure S3.** Flavoaffinin mass spectra. HR-ESI-$MS^1$ spectra of: (A) Flavoaffinin 423, (B) Flavoaffinin 449, (C) Flavoaffinin 407, and (D) Flavoaffinin 433. As noted in the legend of Figure 3 (*Main text*), polyenes often show ionization by in-source dehydrogenation (resulting in an $[M-H]^+$ peak) and ionization as the radical cation (resulting in an $[M]^+$ peak). These two additional peaks are clearly visible in all $MS^1$ spectra except that of flavoaffinin 407.

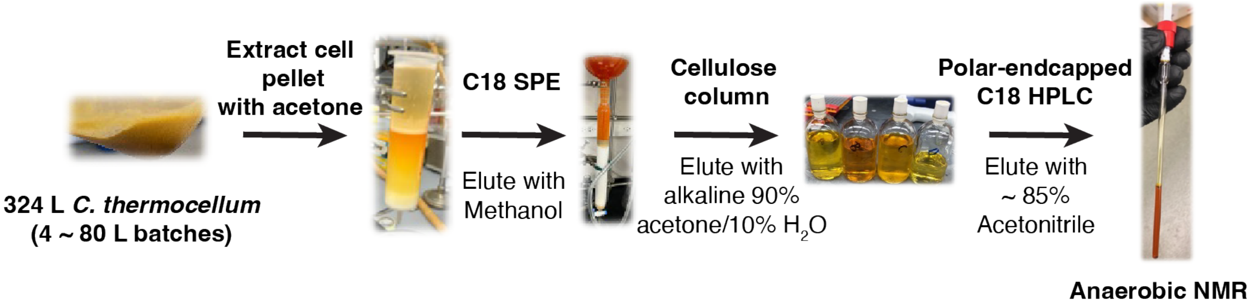

**Figure S4**. Flavoaffinin purification process. The purification could be conducted under an ambient atmosphere, but exposure to light was prevented whenever possible. Even in the dark and at 4 °C, the preparations lost color under ambient atmosphere over the course of a few days. Anoxic NMR in a vacuum tube was essential for compound stability over the course of data acquisition.

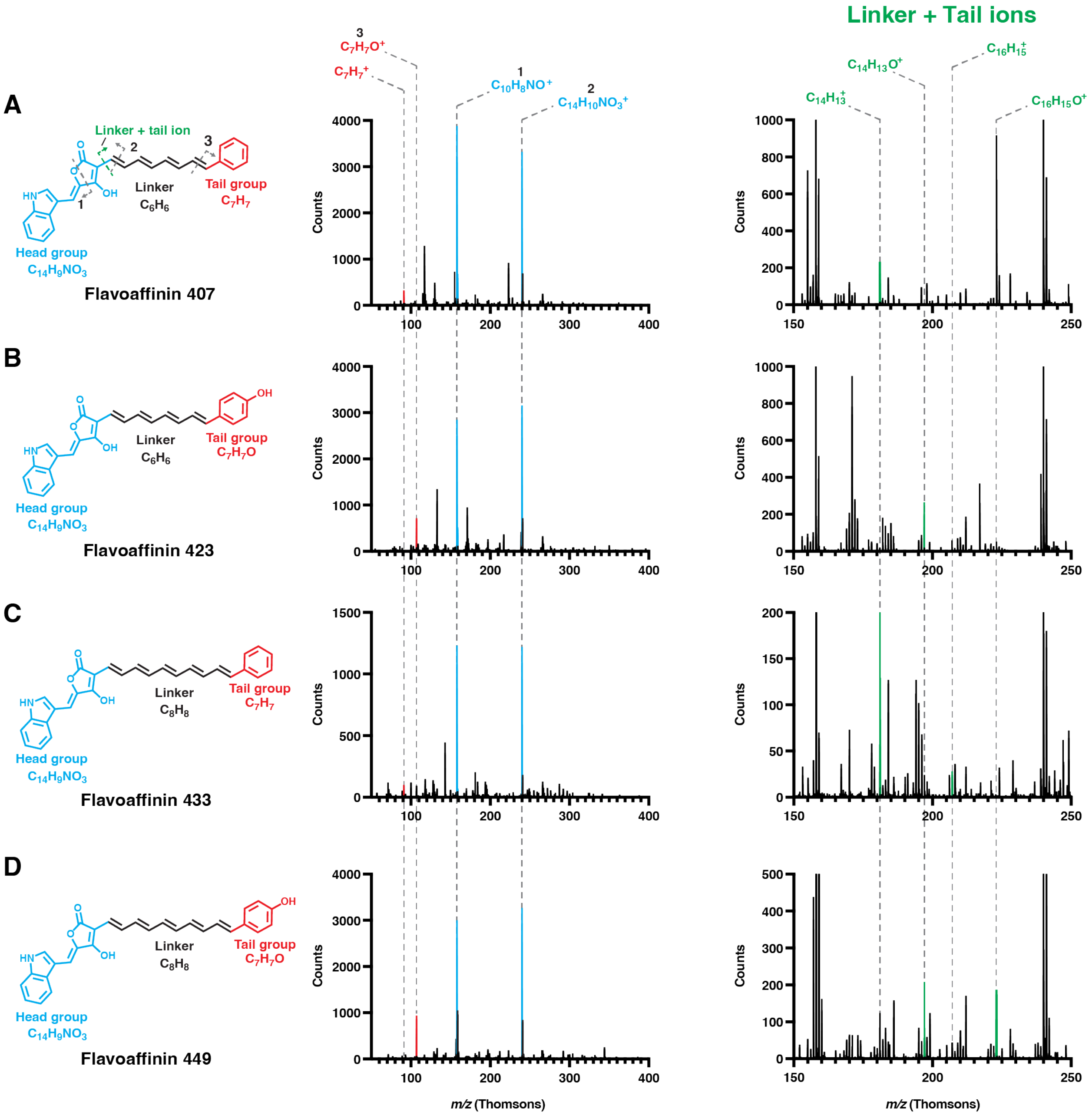

**Figure S5.** Comparison of the HR-ESI-MS[2] spectra of the four flavoaffinin congeners from *C. thermocellum*. (A) Flavoaffinin 407. (B) Flavoaffinin 423. (C) Flavoaffinin 433. (D) Flavoaffinin 449. In each panel, the proposed structure for the indicated congener is given on the left, the MS[2] spectrum for that congener in the *m/z* 50–400 region (showing “head” and “tail” group fragment ions) is given in the center, and a magnified view of the corresponding *m/z* 150–250 region (showing “linker + tail” group ions) is given on the right. In each panel, components of the conserved “head” group and its fragment ions are colored blue, the components of the “tail” group and its fragments ions are colored red, and the ions composed of the tail group and polyene linker are colored green. Vertical dashed lines match ions across panels. For the tail group fragment ions, only the smallest ions containing the entire tail group are shown. For the “linker + tail” ions, only the largest such fragment ions are shown, except in cases were the “linker + tail” ion is seen in another panel.

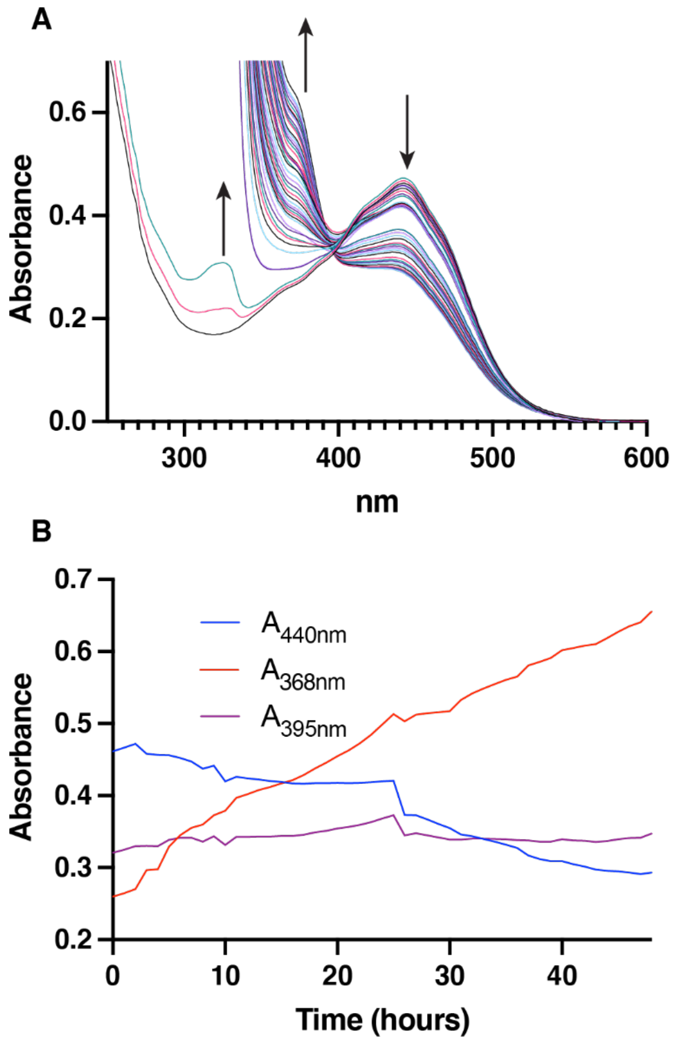

**Figure S6.** Degradation of the flavoaffinins in air. An acetone extract of residual cellulose from a *C. thermocellum* culture was diluted 1:9 in methanol and the UV-visible spectrum was followed with a Cary 3500 Multicell UV-Vis Spectrophotometer (Agilent) during incubation in the dark under ambient atmosphere and at 25 °C. (A) The UV-visible spectrum of the diluted extract over 48 hours with measurements taken every hour. (B) Absorbance traces of the approximate $\lambda_{max}$ of the flavoaffinin absorption band (440 nm) as well as at 368 nm and 395 nm. The $\lambda_{max}$ = 368 nm absorbance band seems to correspond to the $\lambda_{max}$ = 375 nm absorbance band observed by Ljungdahl *et al.*[4] upon storage of crude flavoaffinin preparations under air for 24 hours (spectrum taken in acetone). The 395 nm trace corresponds to an approximate isosbestic point for the UV-visible spectra, especially after 26 hours of incubation.

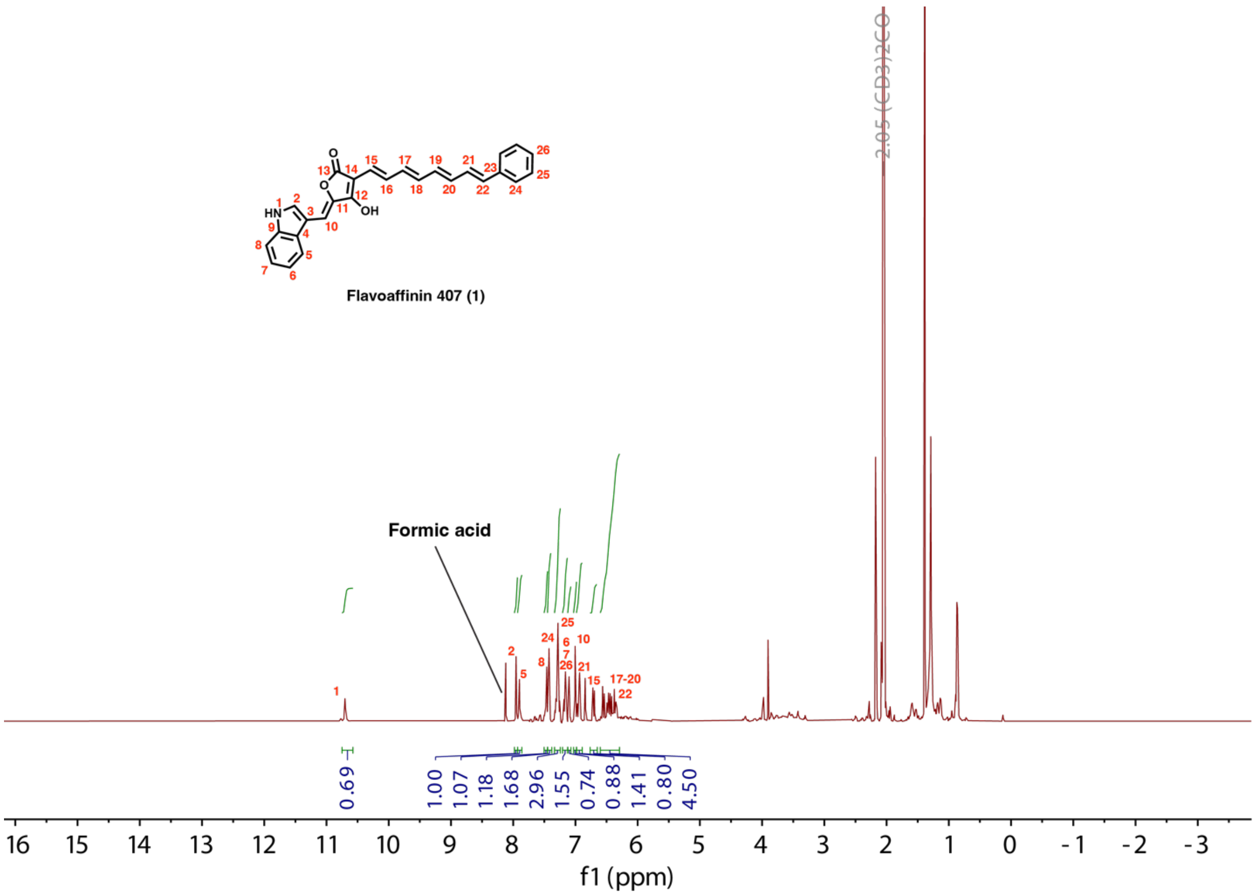

**Figure S7.** [1]H NMR (600 MHz) spectrum of flavoaffinin 407 in $(CD_3)_2CO$.

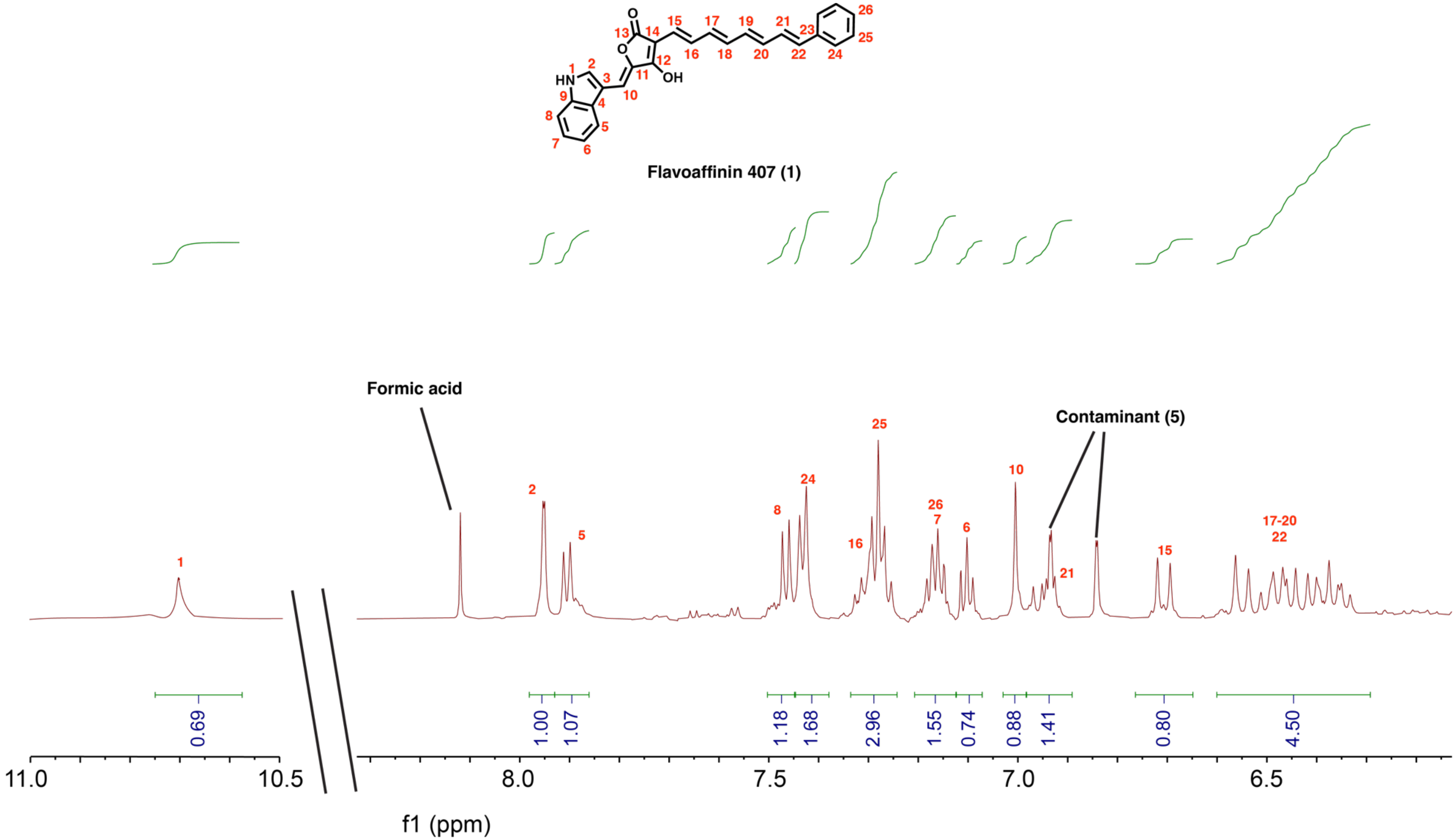

**Figure S8.** $^1$H NMR (600 MHz) spectrum of flavoaffinin 407 in $(CD_3)_2CO$ (view of only the resonances assigned to flavoaffinin 407).

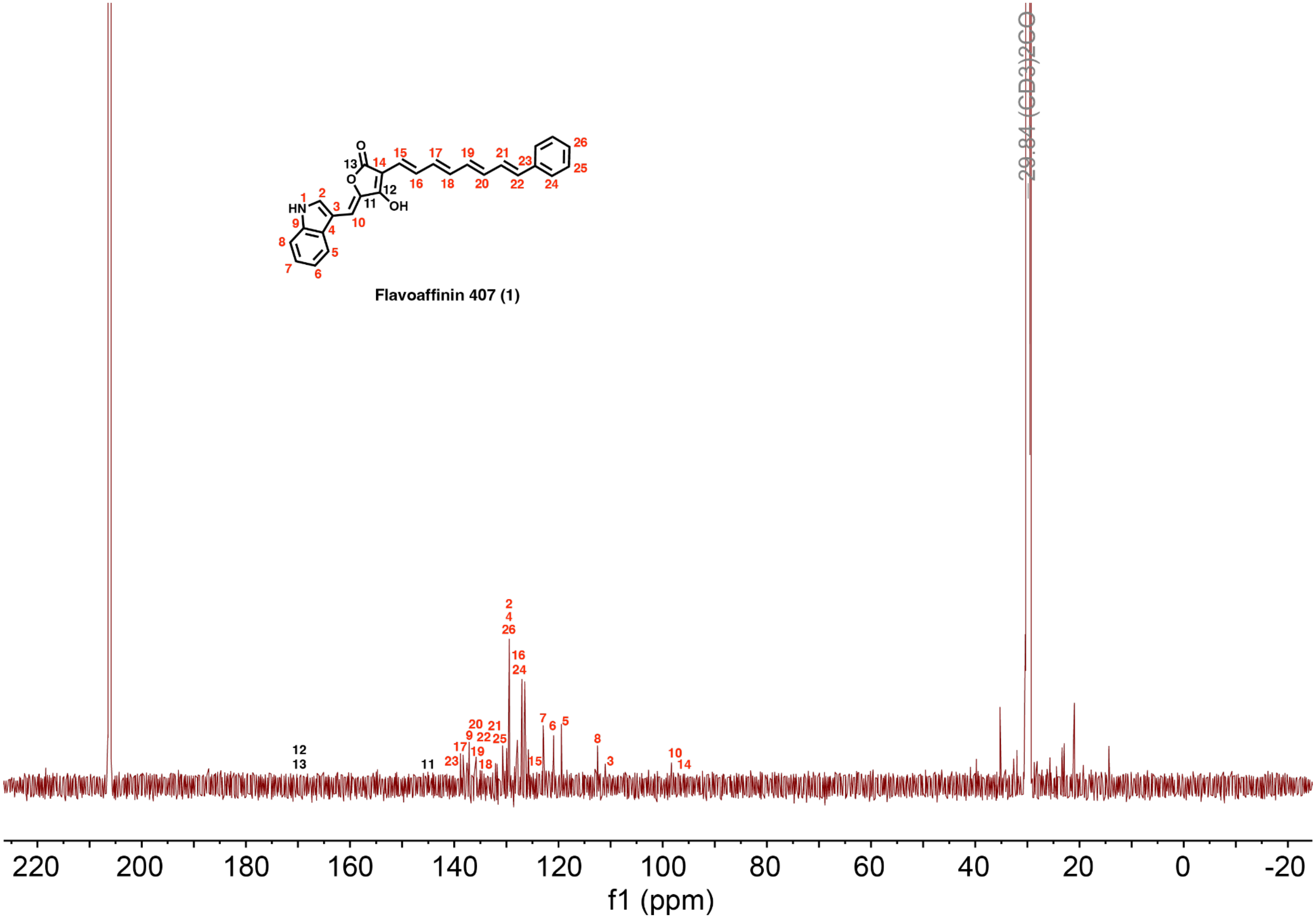

**Figure S9.** $^{13}C$ NMR spectrum (125 MHz) of flavoaffinin 407 in $(CD_3)_2CO$. Position numbers given in black (11-13) correspond to resonances that were not sufficiently intense to be distinguished from background noise.

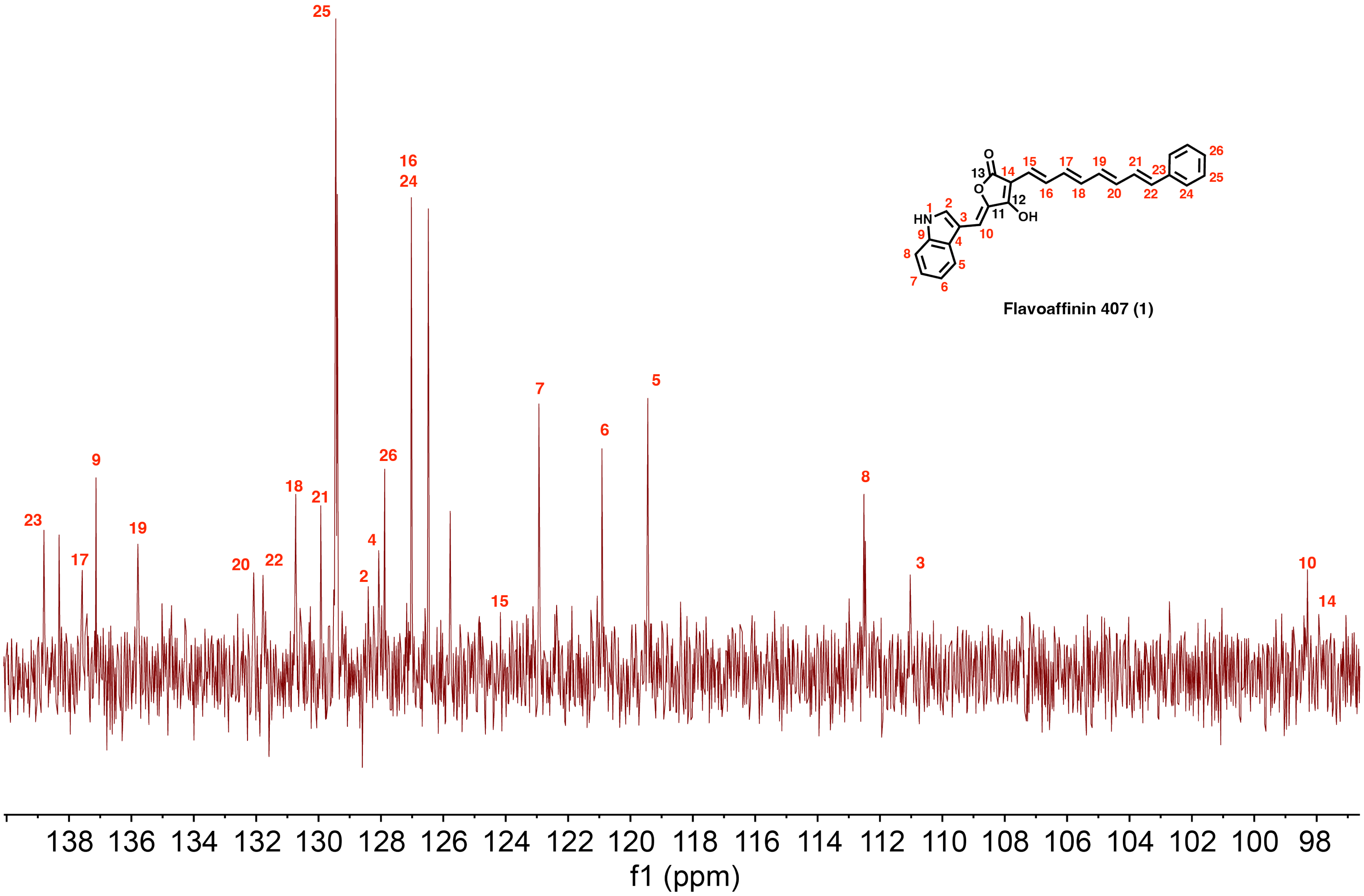

**Figure S10.** $^{13}C$ NMR spectrum (125 MHz) of flavoaffinin 407 in $(CD_3)_2CO$ (view of only the resonances assigned to flavoaffinin 407). Position numbers given in black (11-13) correspond to resonances that were not sufficiently intense to be distinguished from background noise. They are not shown in the spectrum as they are downfield of all other resonances in the molecule.

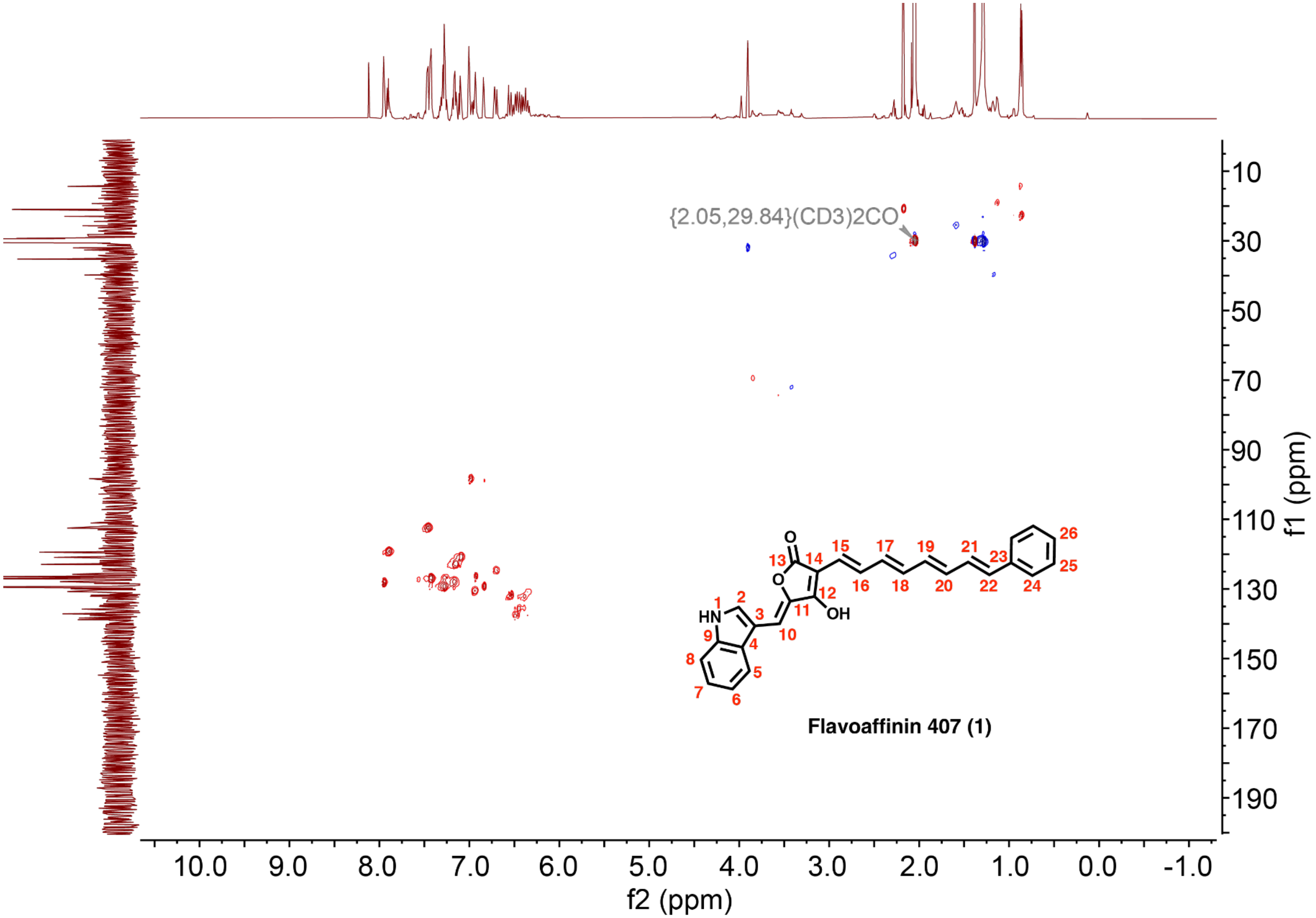

**Figure S11.** HSQC spectrum of flavoaffinin 407 in $(CD_3)_2CO$.

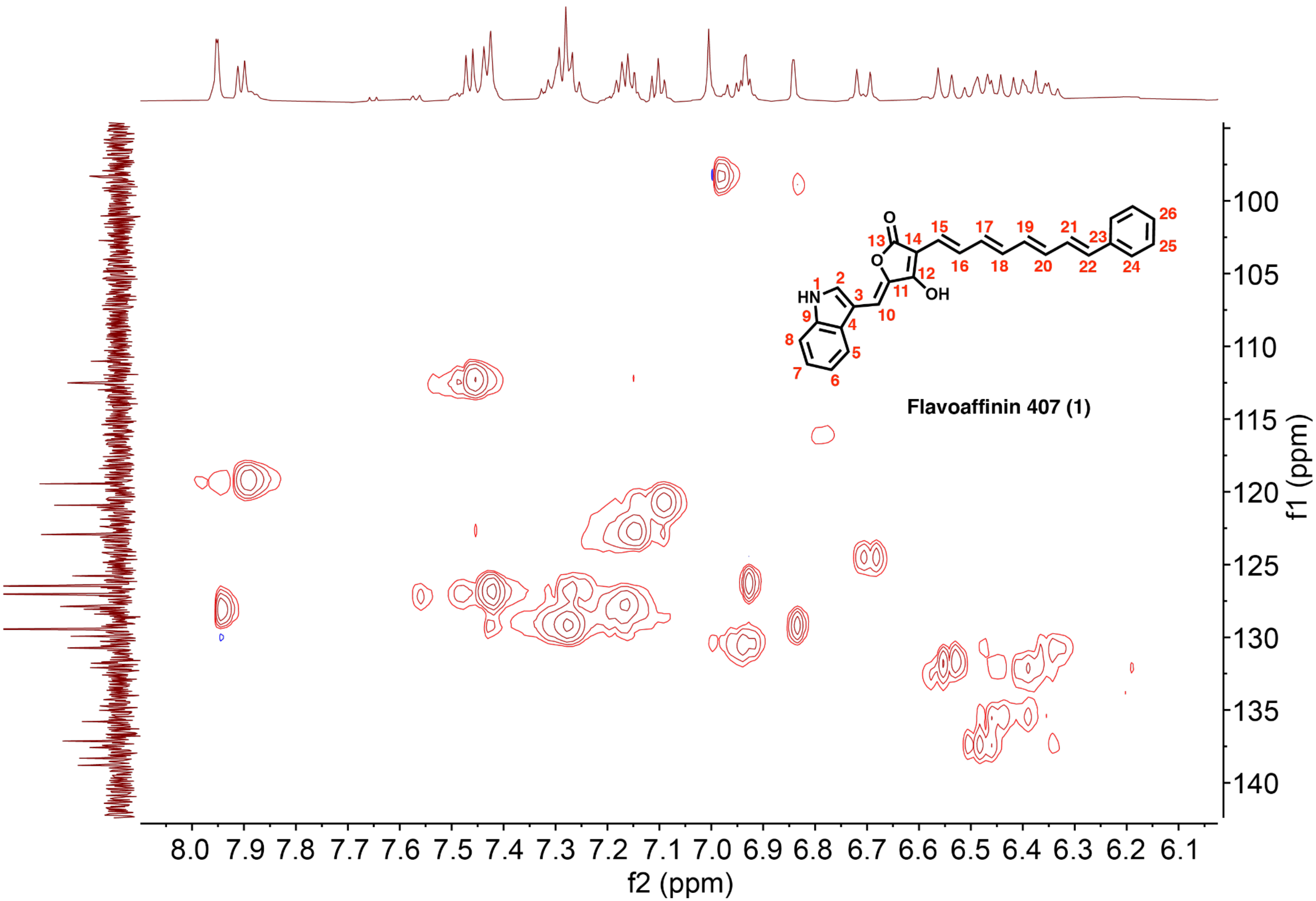

**Figure S12.** HSQC spectrum of flavoaffinin 407 in $(CD_3)_2CO$ (view of only the resonances assigned to flavoaffinin 407).

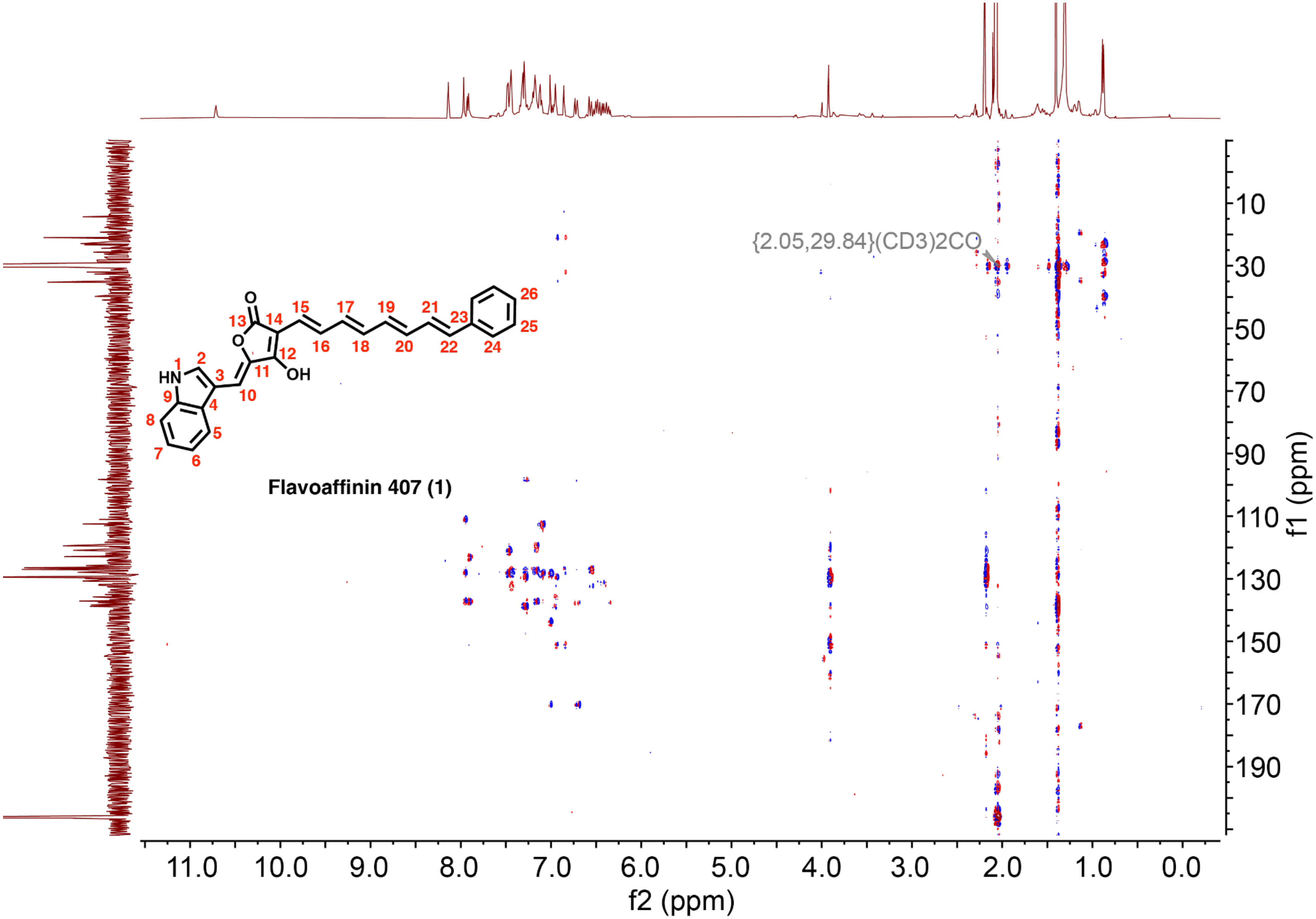

**Figure S13.** HMBC spectrum of flavoaffinin 407 in $(CD_3)_2CO$.

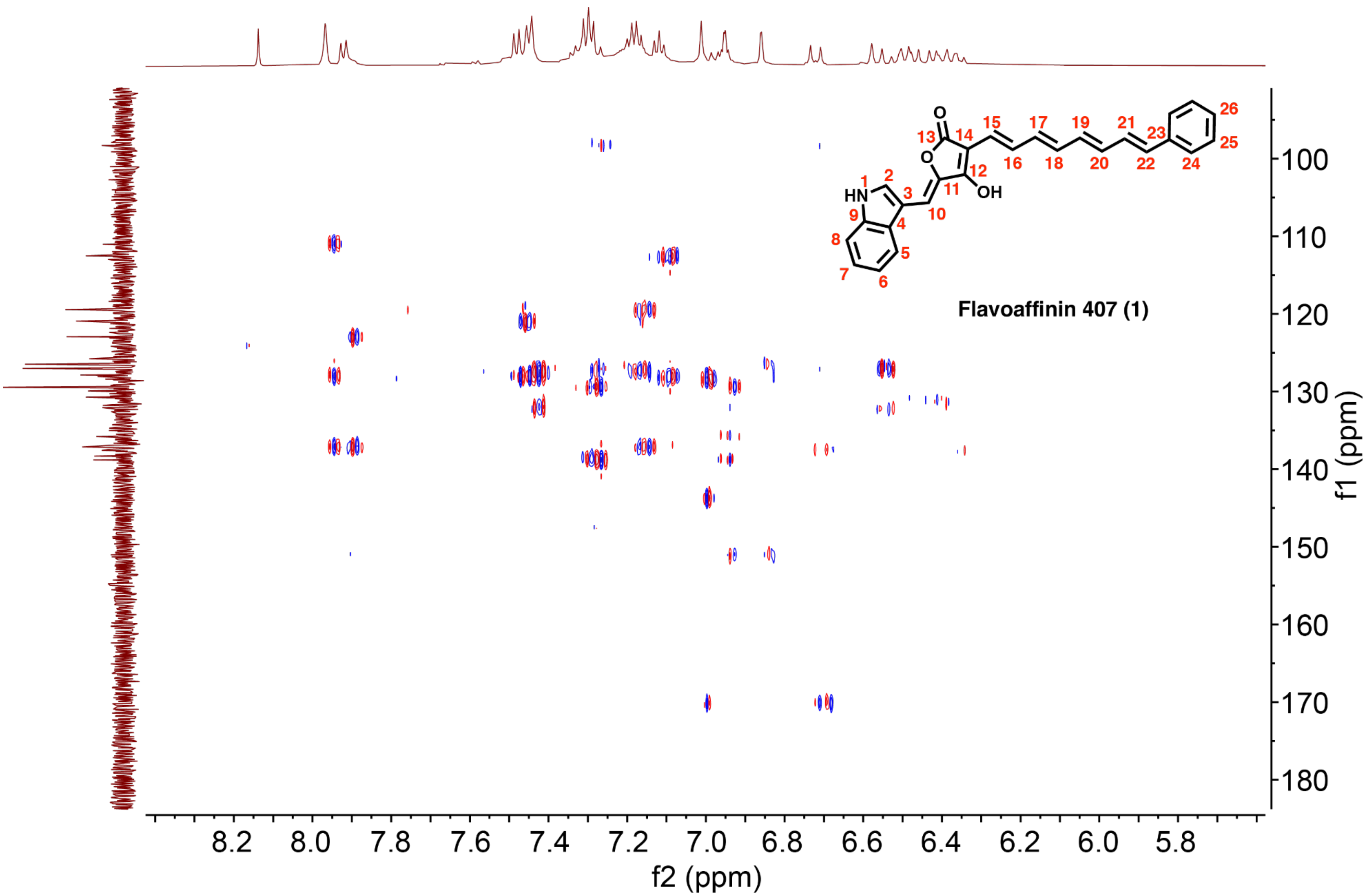

**Figure S14.** HMBC spectrum of flavoaffinin 407 in $(CD_3)_2CO$ (view of only the resonances assigned to flavoaffinin 407).

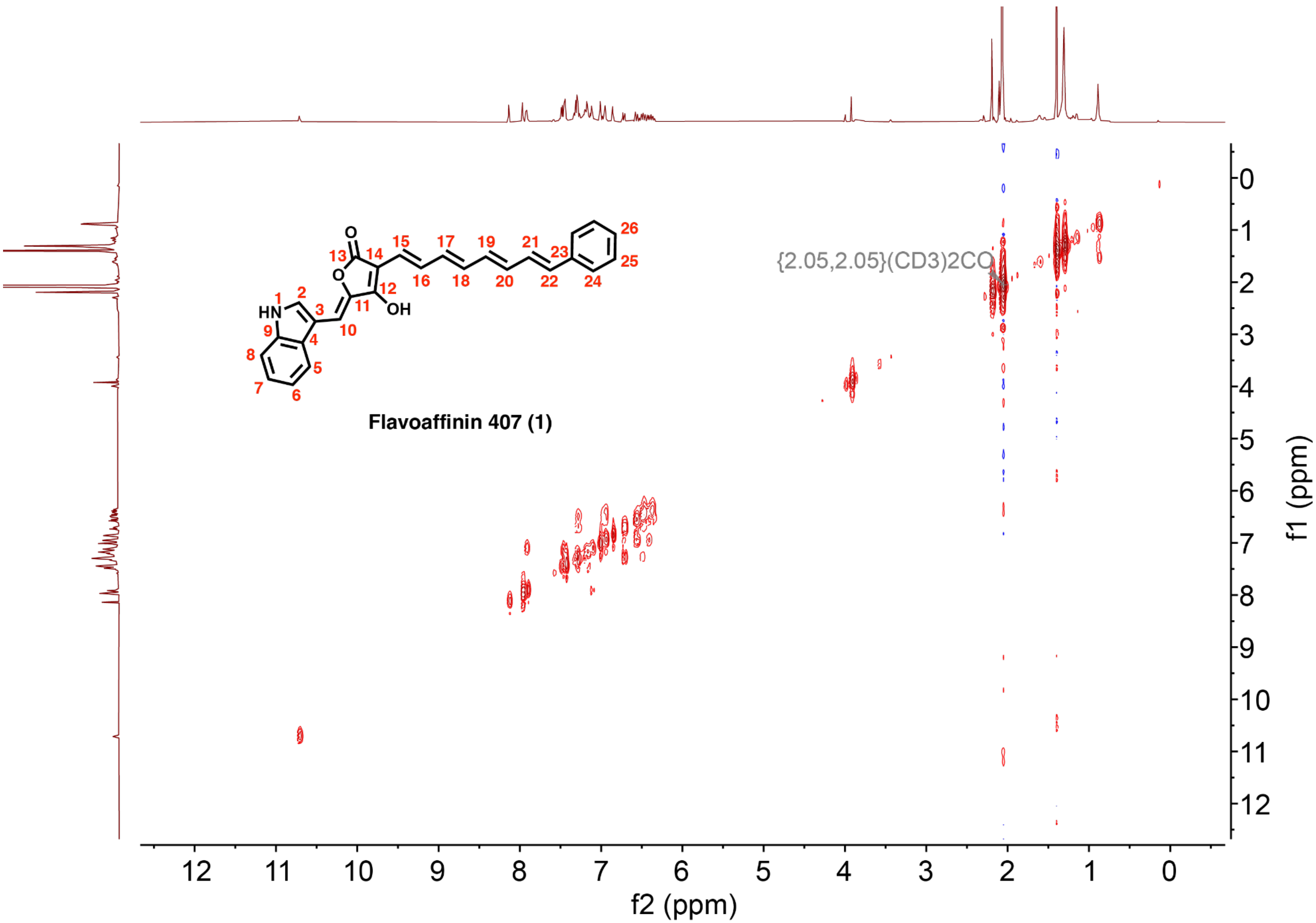

**Figure S15.** $^1$H-$^1$H COSY spectrum of flavoaffinin 407 in $(CD_3)_2CO$.

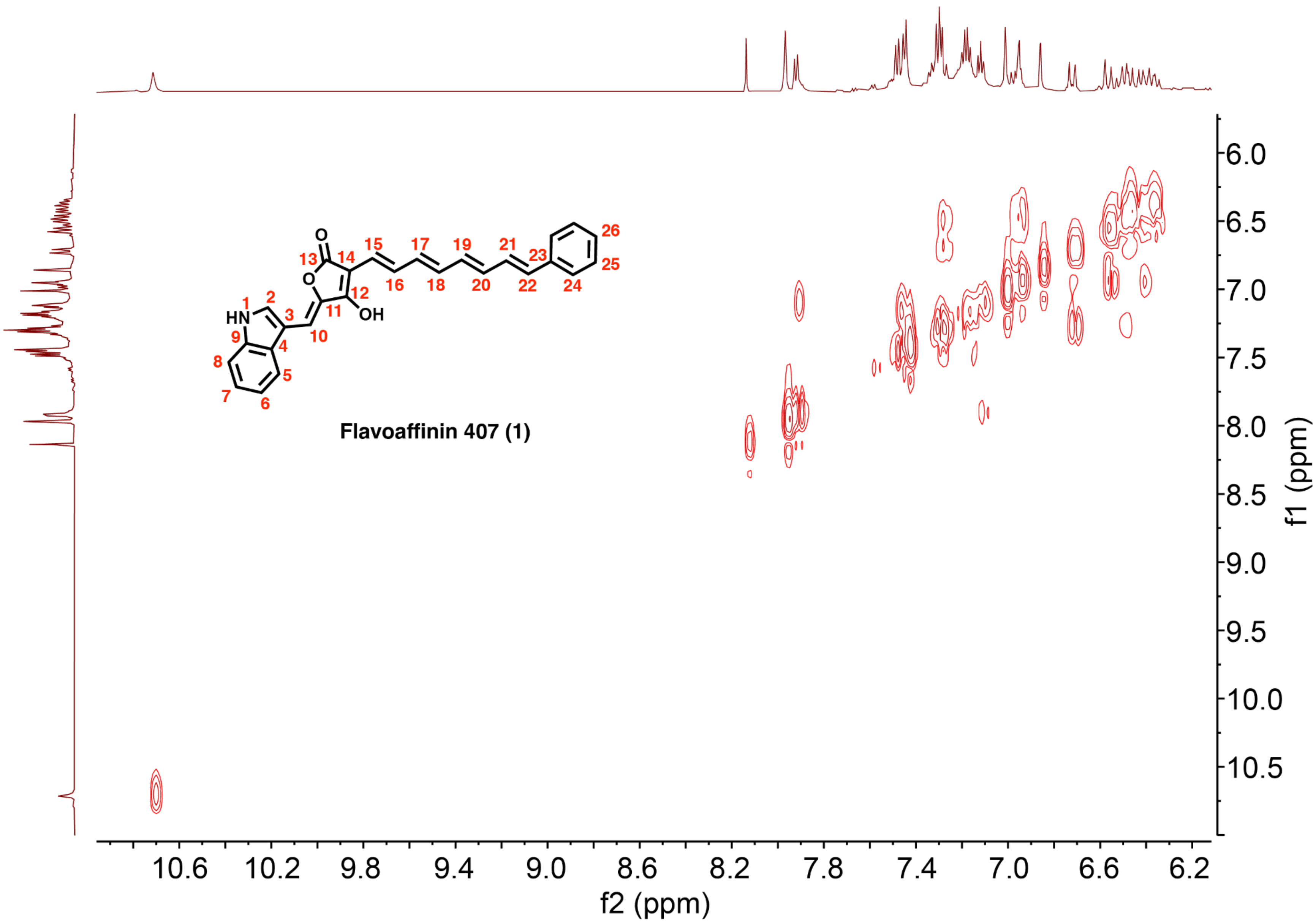

**Figure S16.** $^1H$-$^1H$ COSY spectrum of flavoaffinin 407 in $(CD_3)_2CO$ (view of only the resonances assigned to flavoaffinin 407).

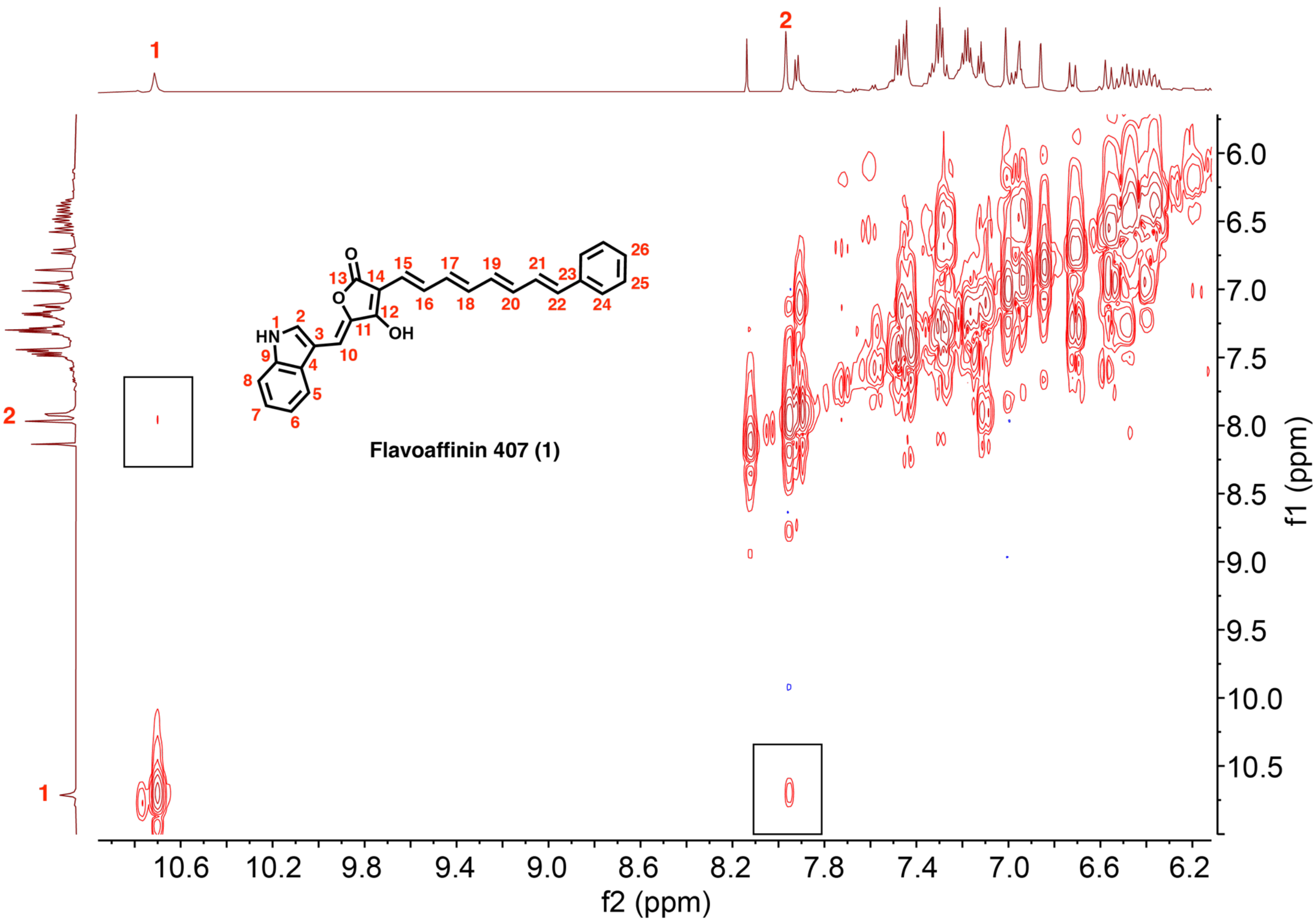

**Figure S17.** [1]H-[1]H COSY spectrum of flavoaffinin 407 in $(CD_3)_2CO$ (view of only the resonances assigned to flavoaffinin 407, and showing weak correlation between the indole N–H1 and H2 in black rectangles).

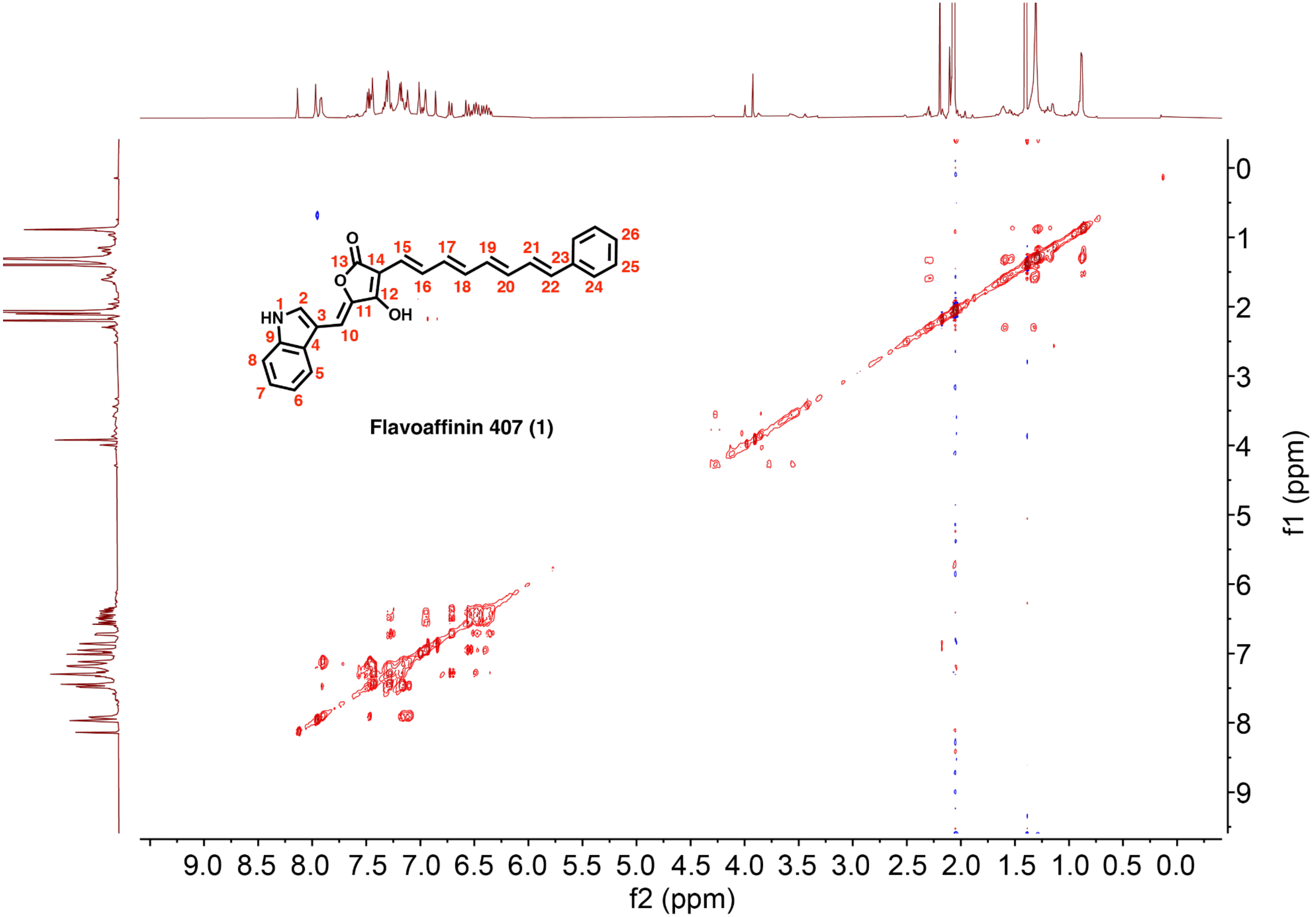

**Figure S18.** TOCSY spectrum of flavoaffinin 407 in $(CD_3)_2CO$. Note that the resonance of H1 is outside of the acquired spectral width.

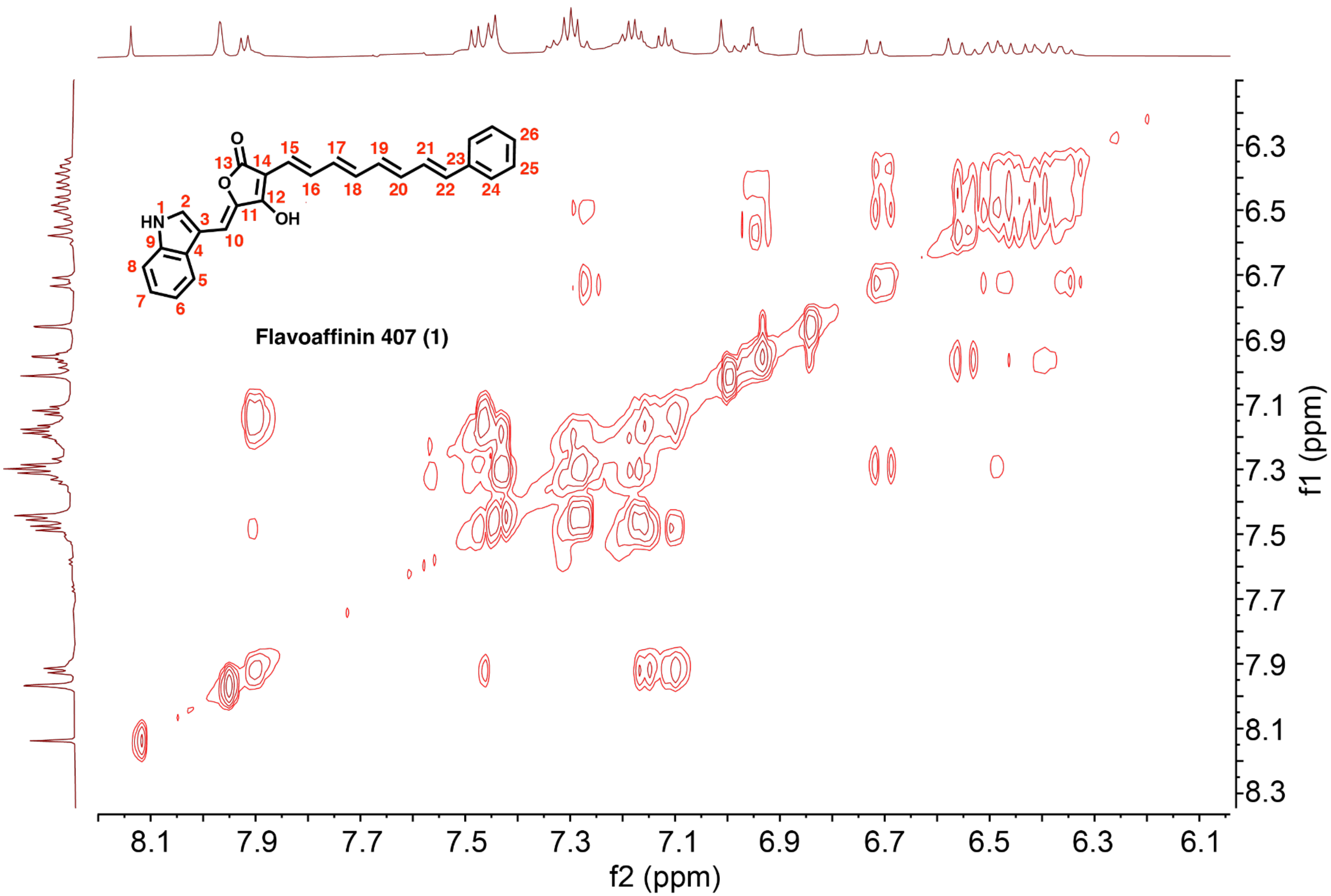

**Figure S19.** TOCSY spectrum of flavoaffinin 407 in $(CD_3)_2CO$ (view of only the resonances assigned to flavoaffinin 407). Note that the resonance of H1 is outside of the acquired spectral width.

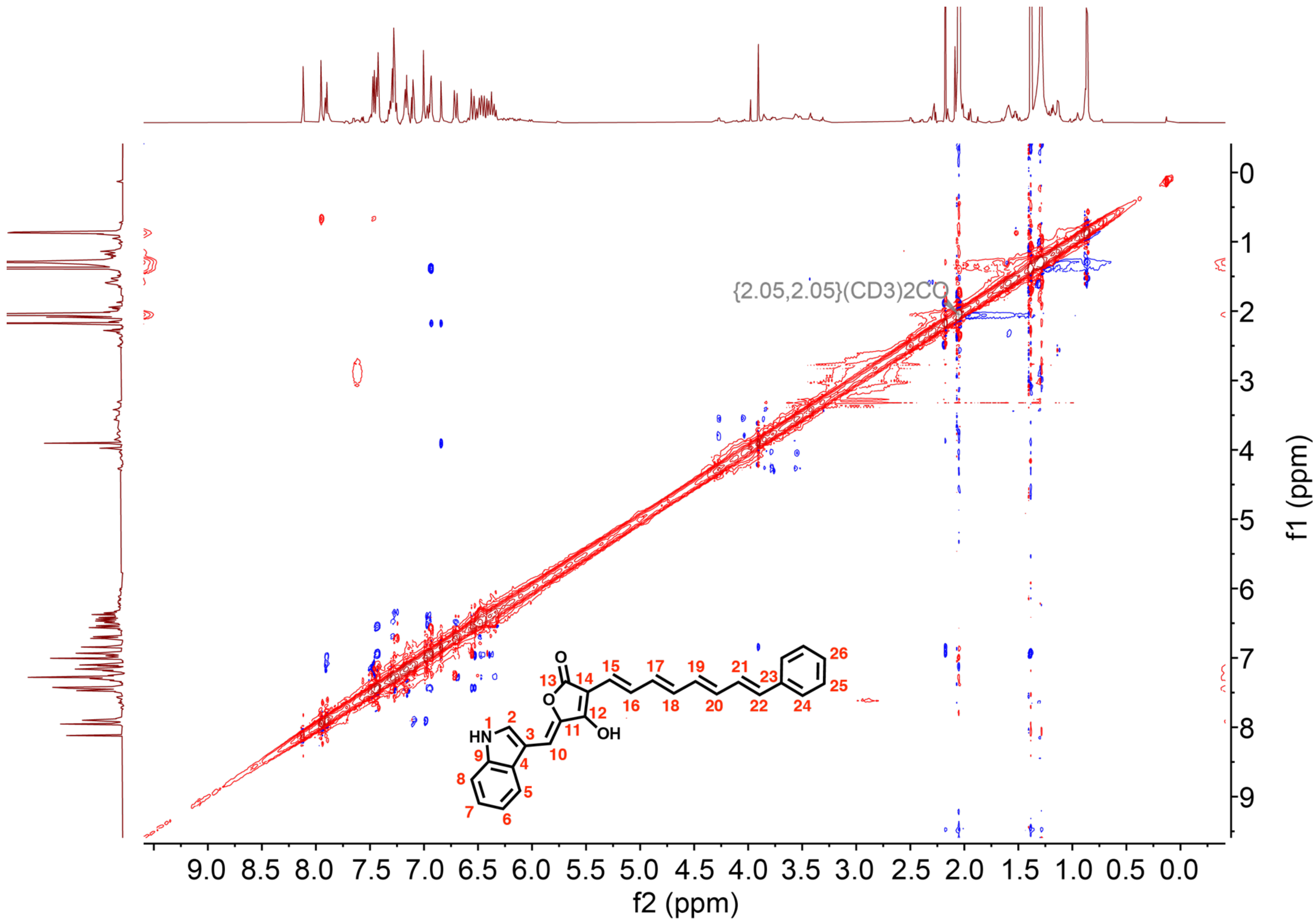

**Figure S20.** NOESY spectrum of flavoaffinin 407 in $(CD_3)_2CO$. Note that the resonance of H1 is outside of the acquired spectral width.

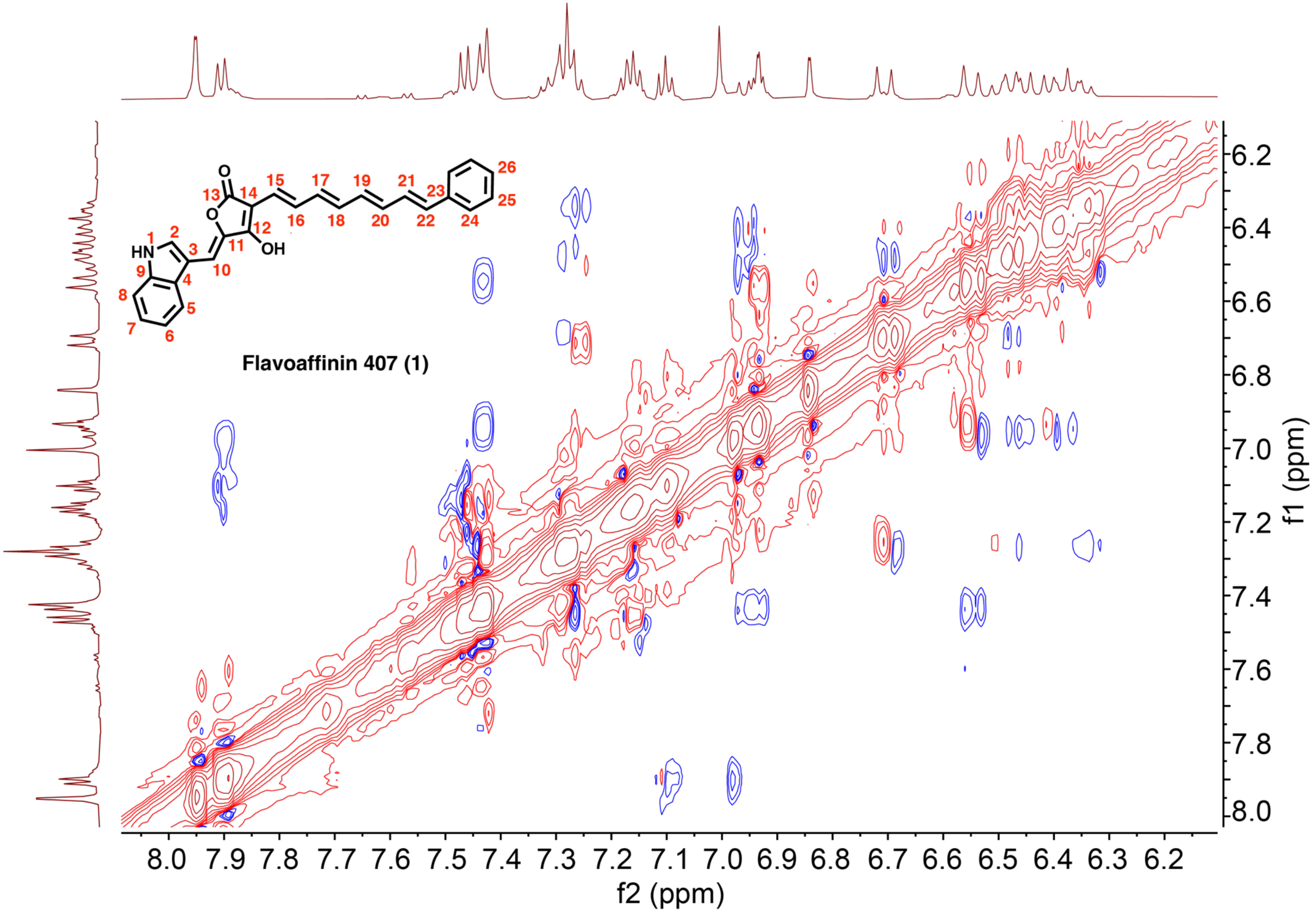

**Figure S21.** NOSEY spectrum of flavoaffinin 407 in $(CD_3)_2CO$ (view of only the resonances assigned to flavoaffinin 407). Note that the resonance of H1 is outside of the acquired spectral width

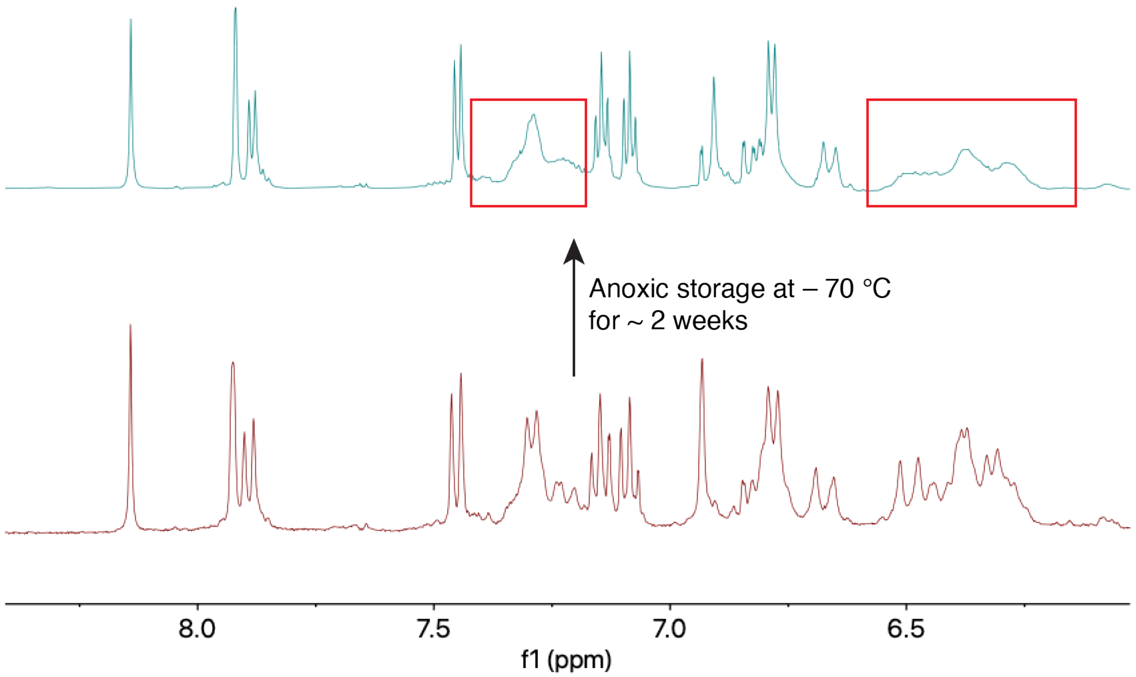

**Figure S22.** Degradation of flavoaffinin 449 observed by $^1$H-NMR. $^1$H-NMR spectra of flavoaffinin 449 in $(CD_3)_2CO$ comparing the freshly prepared sample run on a 400 MHz instrument (bottom, maroon) to the same sample run on a 600 MHz instrument following anoxic storage in the dark at – 70 °C for about two weeks (top, cyan). Note the apparent loss of resolution of the resonances marked with red boxes in cyan (after storage) spectrum. In the flavoaffinin 407 dataset, the equivalent resonances were assigned to the polyene.

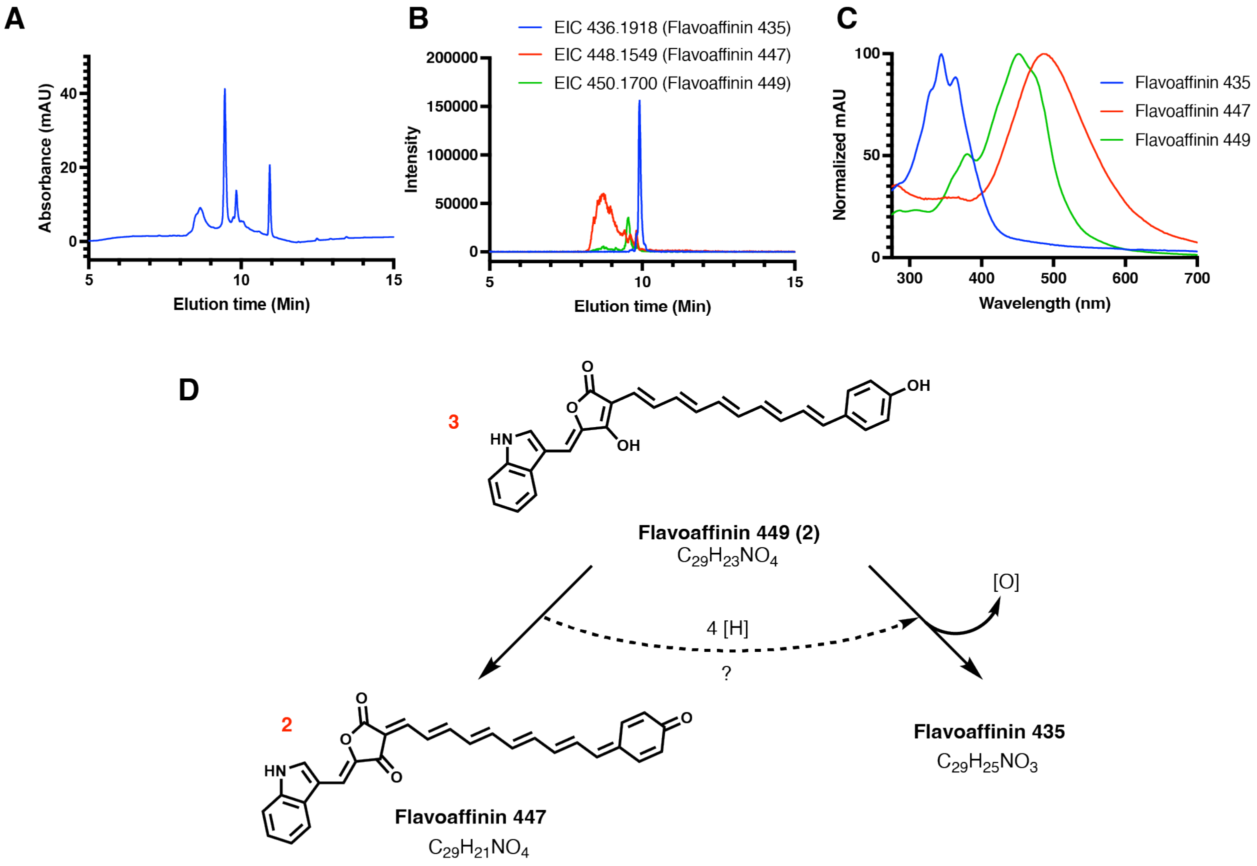

**Figure S23.** Degradation of a purified flavoaffinin 449 sample during anoxic storage at –70 °C. (A) Total wavelength chromatogram (TWC) of a purified flavoaffinin 449 sample (same sample as cyan spectrum in Figure S21) after approximately two weeks of anoxic storage in the dark at – 70 °C. (B) Extracted ion chromatograms (EICs, *m/z* values indicated) of the three identified flavoaffinin metabolites (flavoaffinins 435, 447, and 449) from the sample from Panel A. (C) UV-Visible spectra of the three flavoaffinin metabolites from Panel B. (D) A hypothetical scheme for degradation of flavoaffinin by electron disproportionation under the conditions of anoxic storage. Note that the hypsochromic shift in the flavoaffinin 449 absorption band to that in flavoaffinin 435 is consistent with interruption of conjugation of the polyene at an unknown site. Red numbers refer to the presumed stoichiometry.

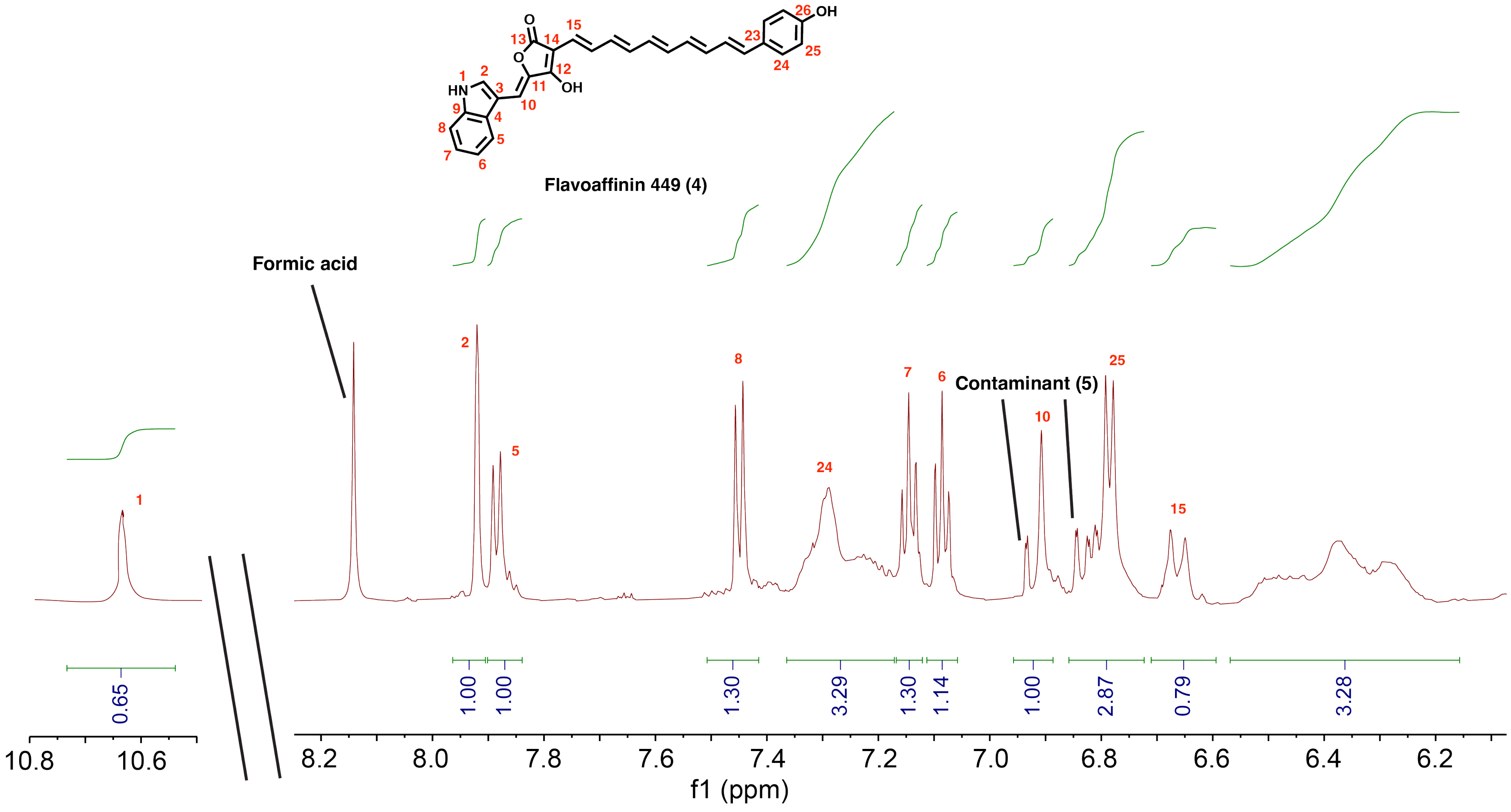

**Figure S24.** $^1$H NMR (600 MHz) spectrum of flavoaffinin 449 in $(CD_3)_2CO$.

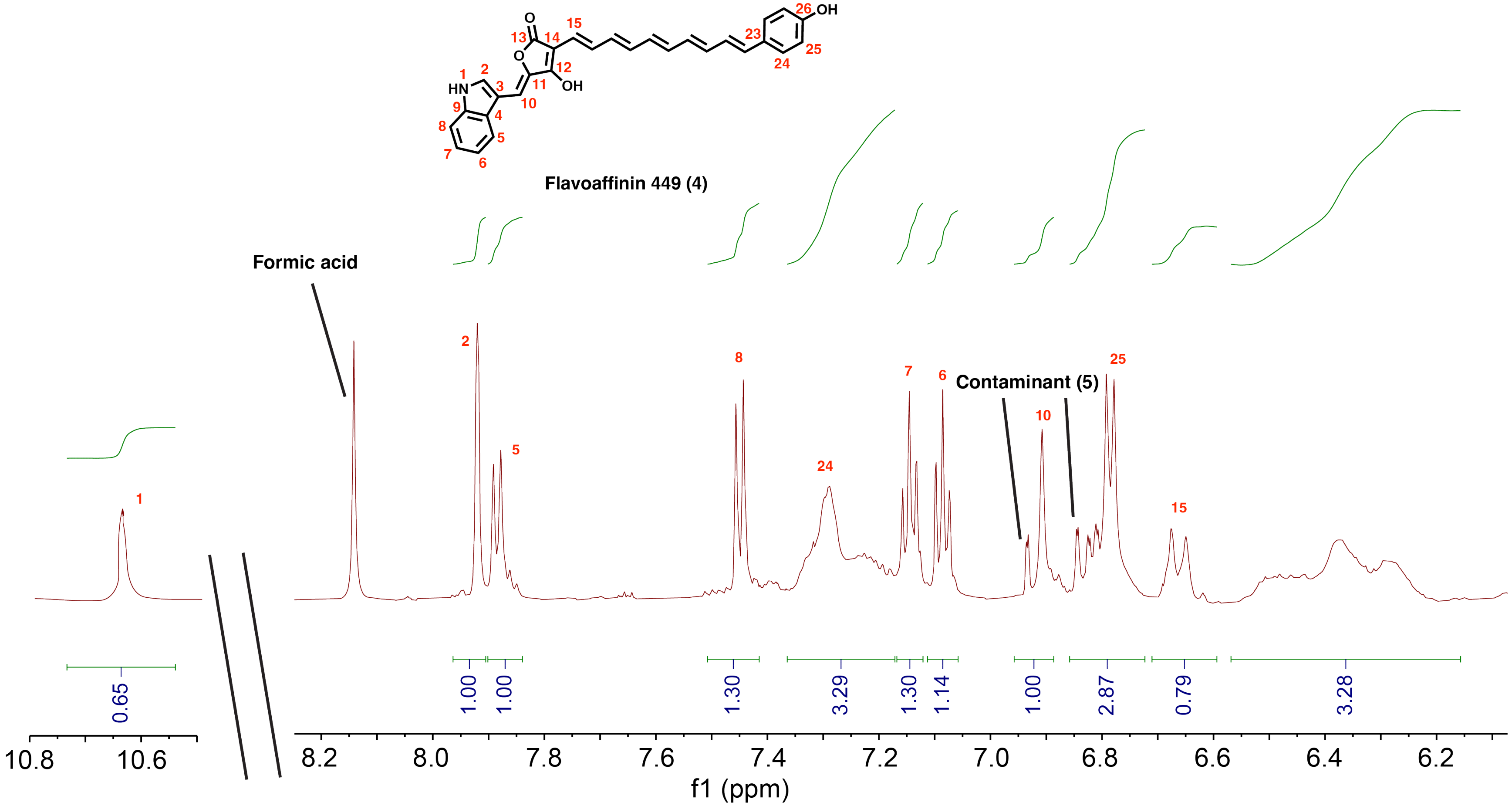

**Figure S25.** $^1H$ NMR (600 MHz) spectrum of flavoaffinin 449 in $(CD_3)_2CO$ (view of only the resonances assigned to flavoaffinin 449).

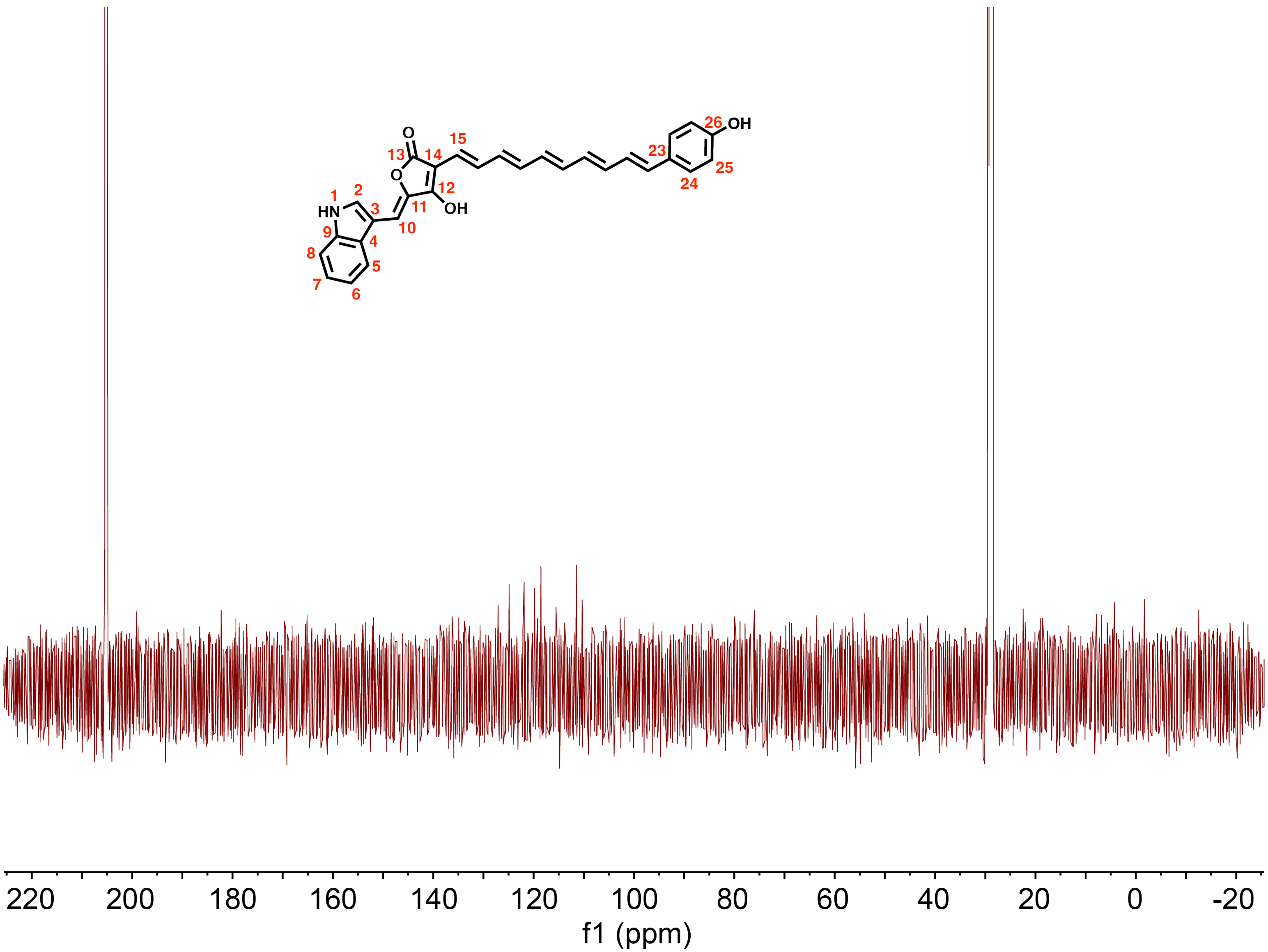

**Figure S26.** $^{13}C$ NMR spectrum (125 MHz) of flavoaffinin 449 in $(CD_3)_2CO$.

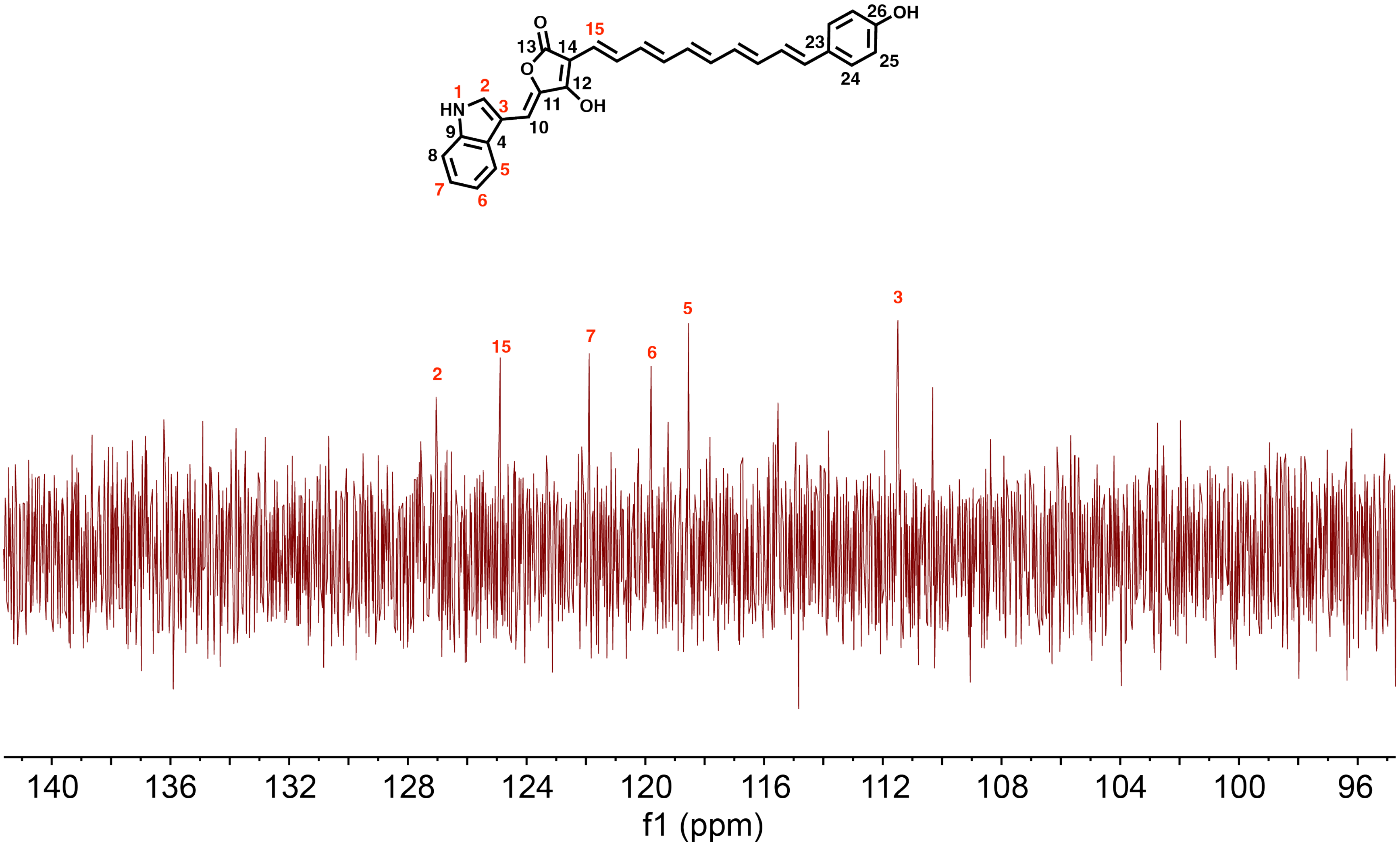

**Figure S27.** $^{13}C$ NMR spectrum (125 MHz) of flavoaffinin 449 in $(CD_3)_2CO$ (view of only the resonances assigned to flavoaffinin 449). Resonances that were judged to be below the limit of detection in this spectrum are numbered in black.

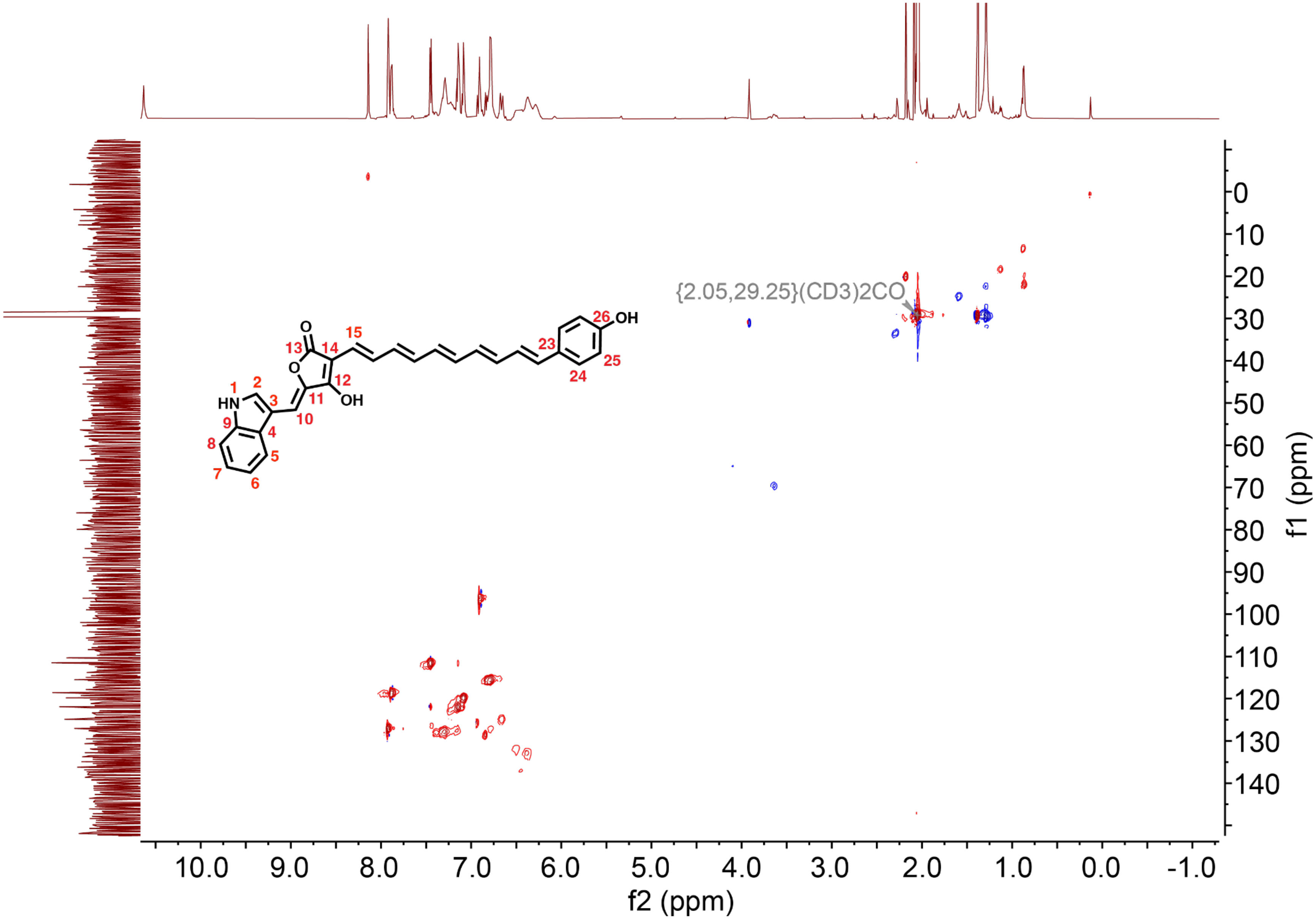

**Figure S28.** HSQC spectrum of flavoaffinin 449 in $(CD_3)_2CO$.

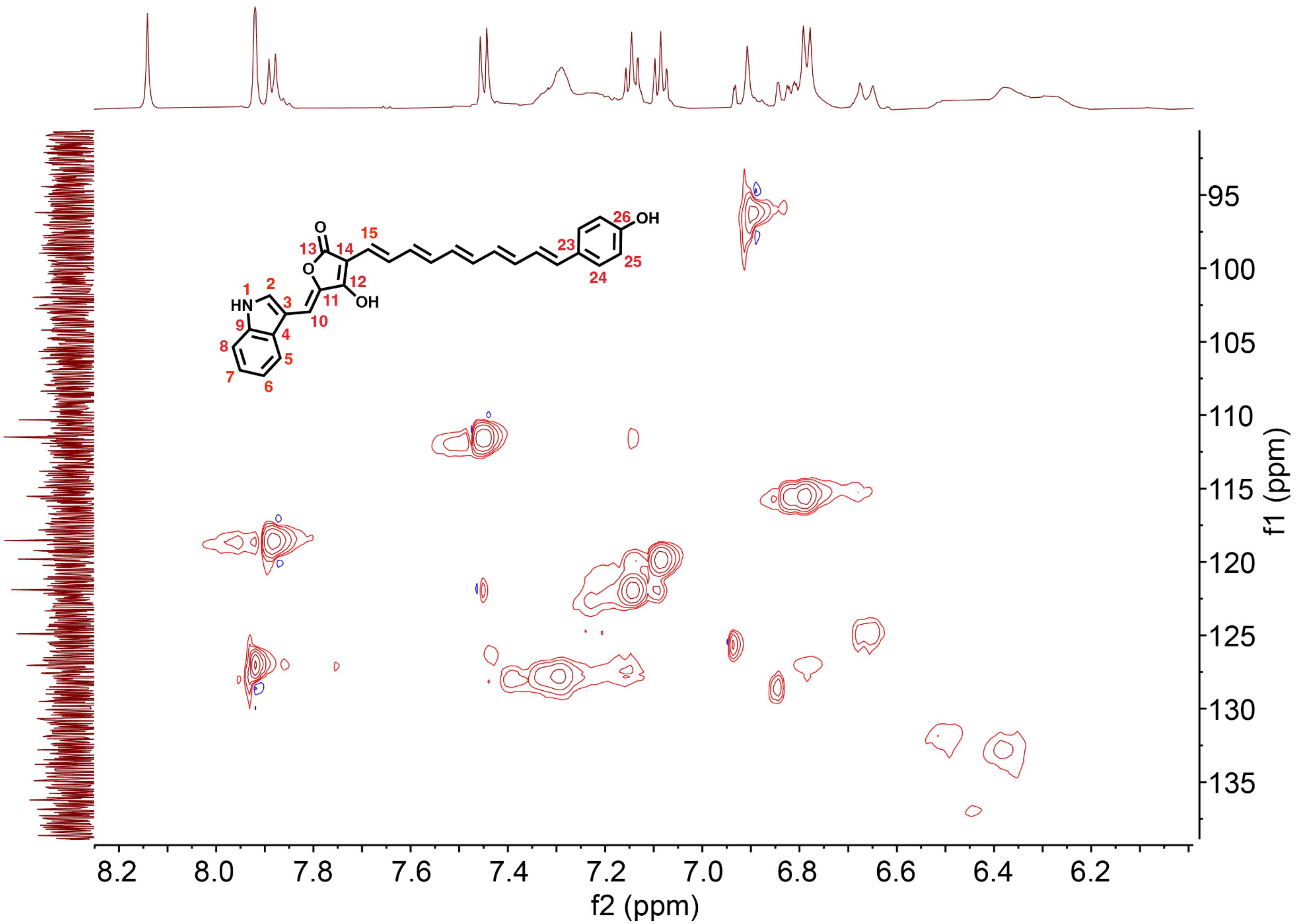

**Figure S29.** HSQC spectrum of flavoaffinin 449 in $(CD_3)_2CO$ (view of only the resonances assigned to flavoaffinin 449).

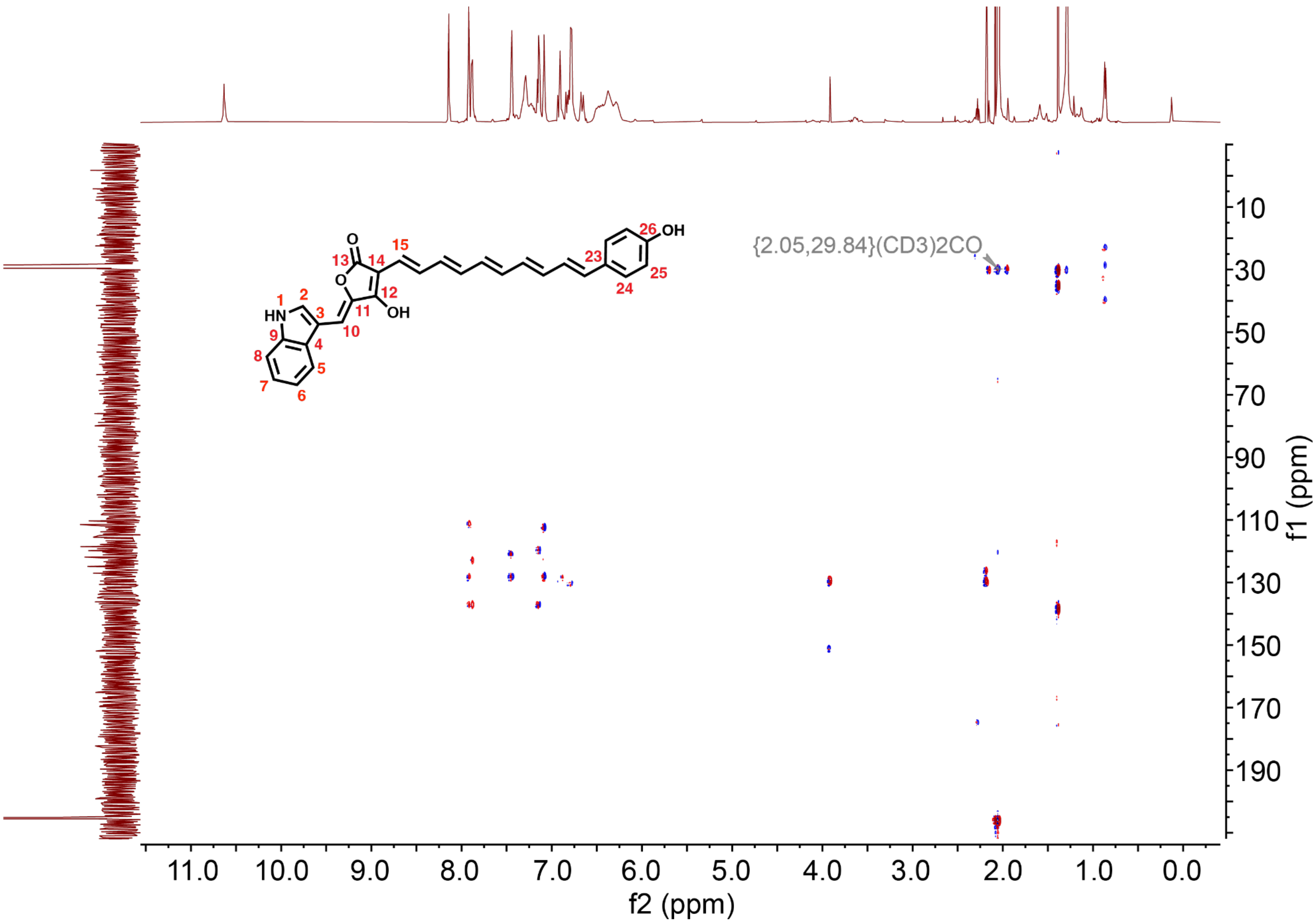

**Figure S30.** HMBC spectrum of flavoaffinin 449 in $(CD_3)_2CO$.

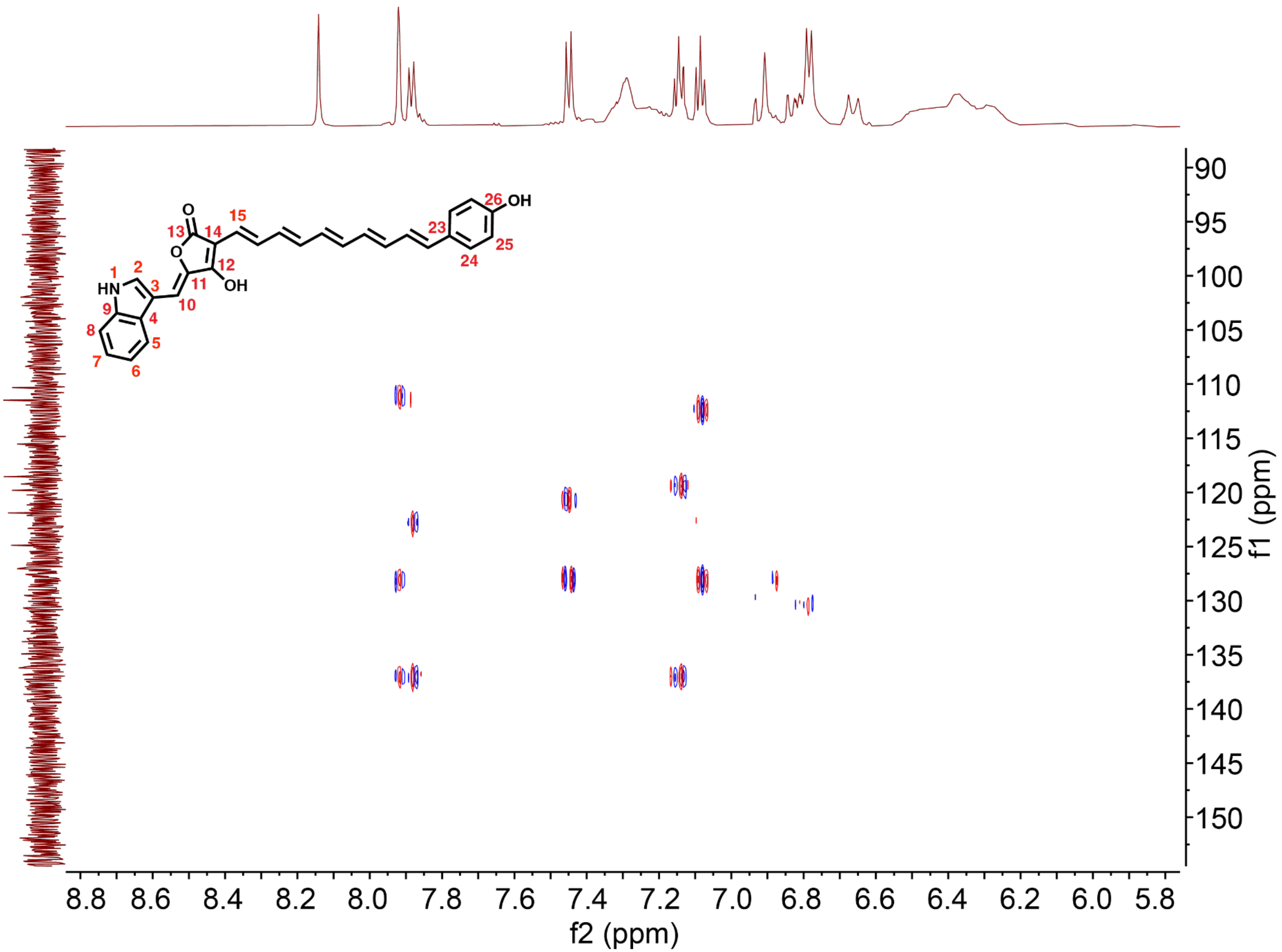

**Figure S31.** HMBC spectrum of flavoaffinin 449 in $(CD_3)_2CO$ (view of only the resonances assigned to flavoaffinin 449).

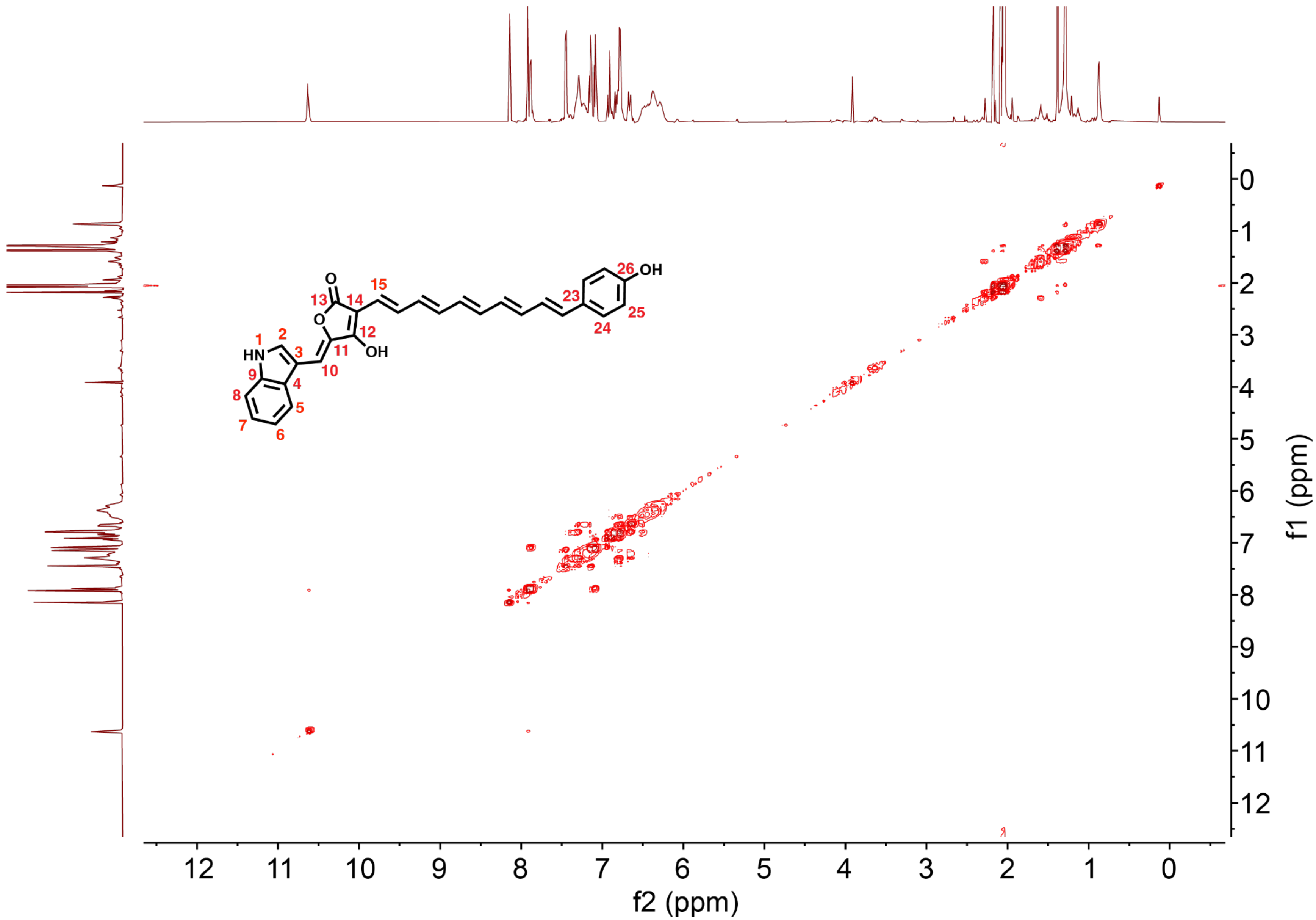

**Figure S32.** $^1$H-$^1$H COSY spectrum of flavoaffinin 449 in $(CD_3)_2CO$.

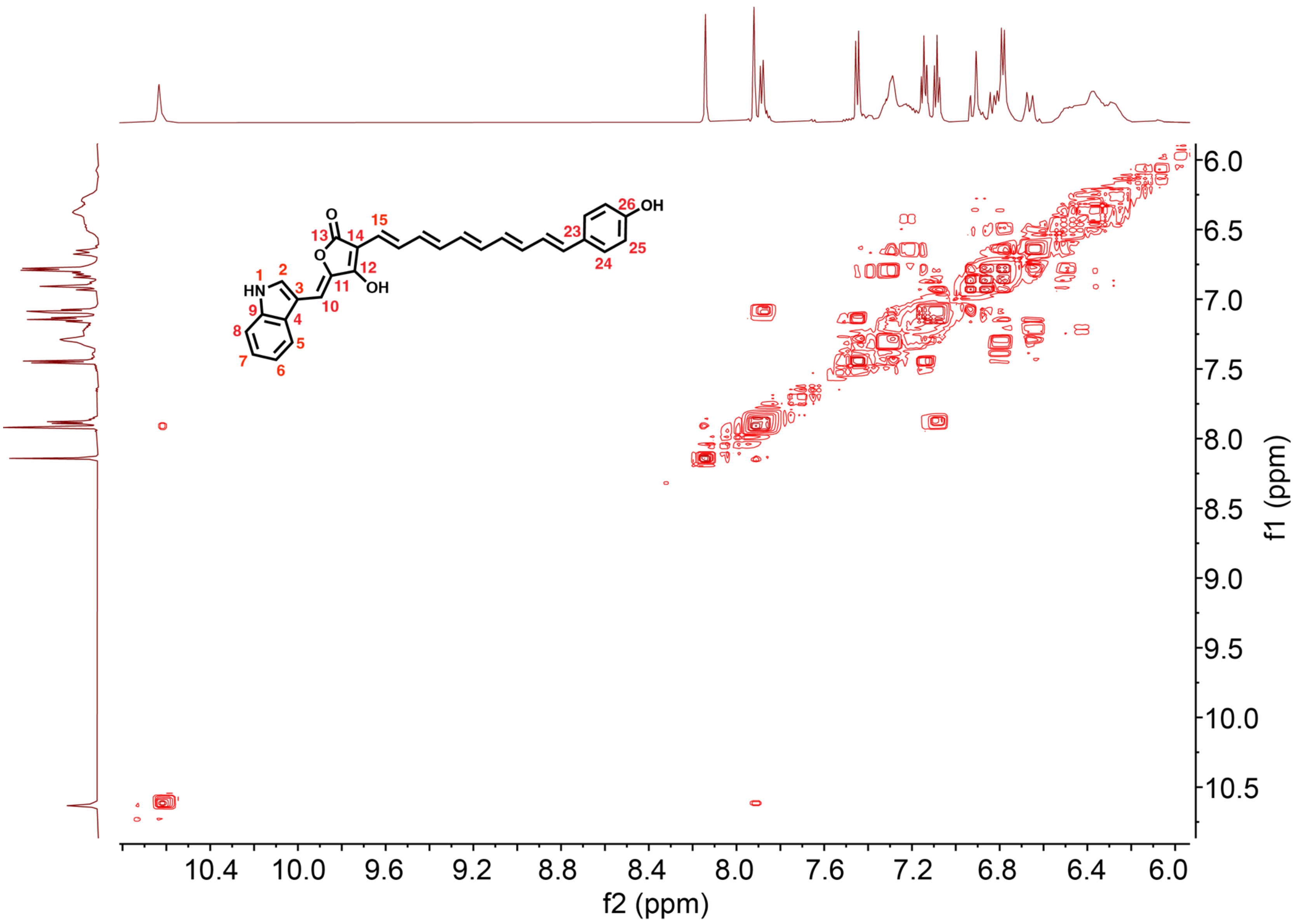

**Figure S33.** [1]H-[1]H COSY spectrum of flavoaffinin 449 in $(CD_3)_2CO$ (view of only the resonances assigned to flavoaffinin 449).

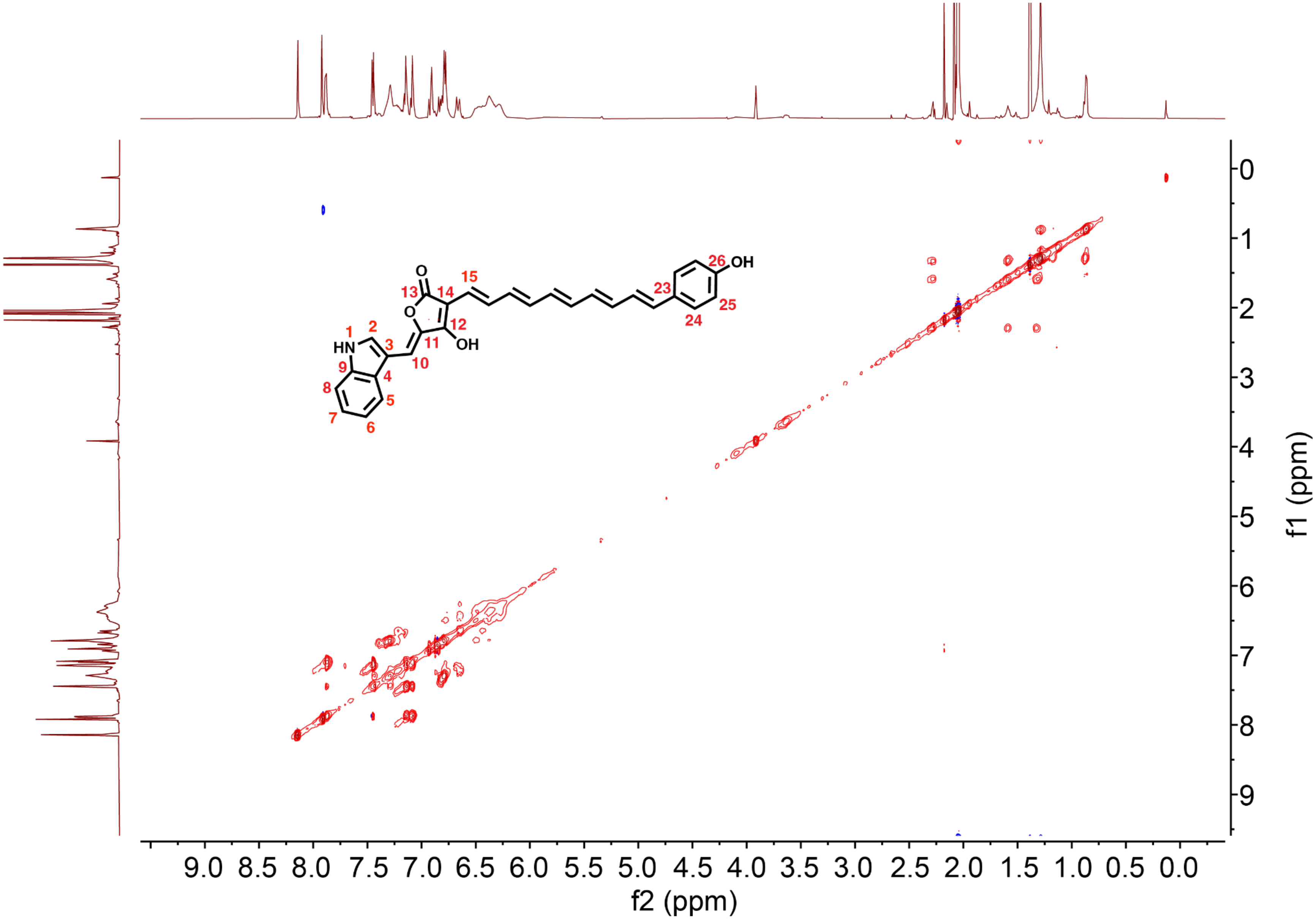

**Figure S34.** $^1H$-$^1H$ TOCSY spectrum of flavoaffinin 449 in $(CD_3)_2CO$.

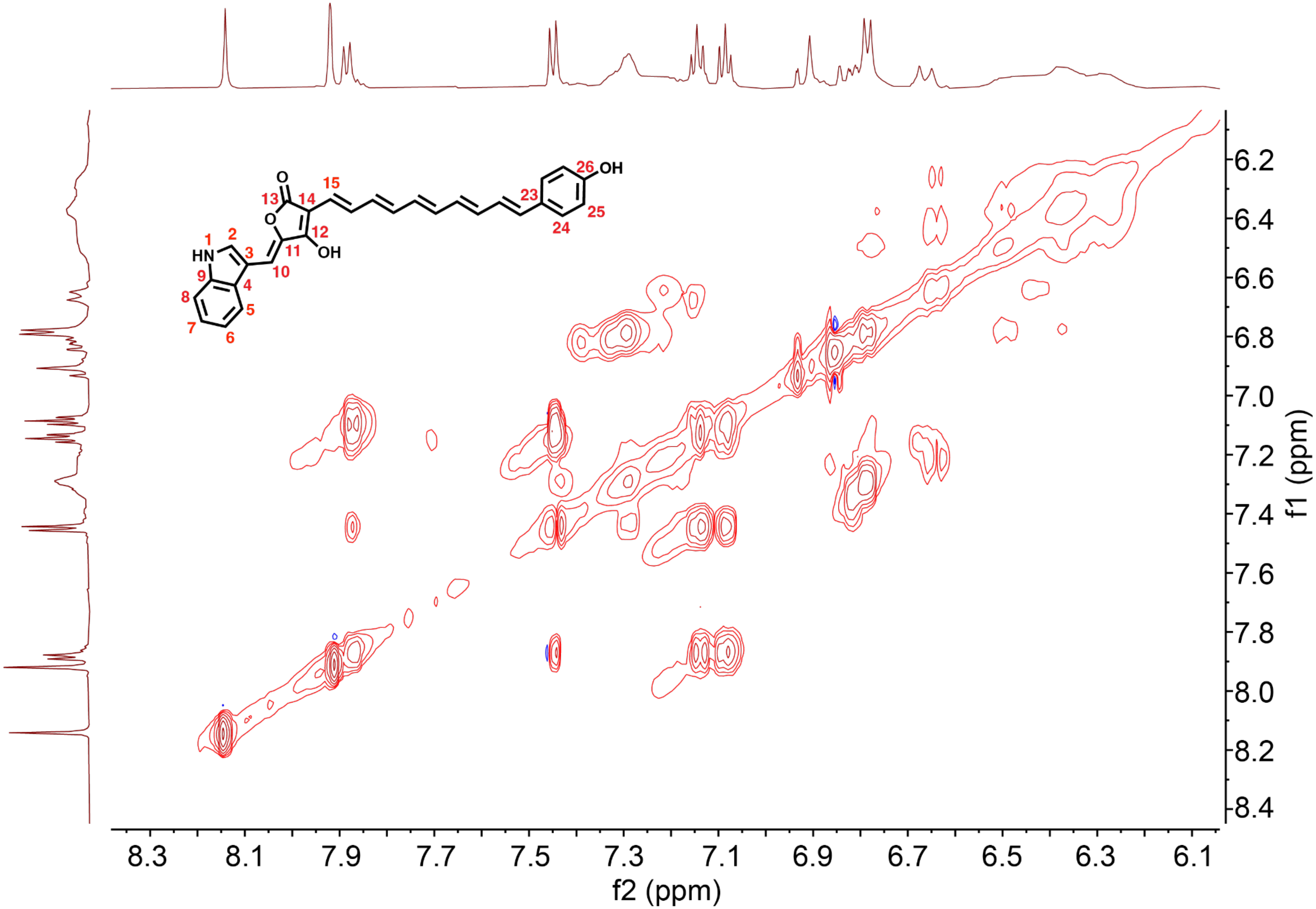

**Figure S35.** $^{1}H$-$^{1}H$ TOCSY spectrum of flavoaffinin 449 in $(CD_3)_2CO$ (view of only the resonances assigned to flavoaffinin 449).

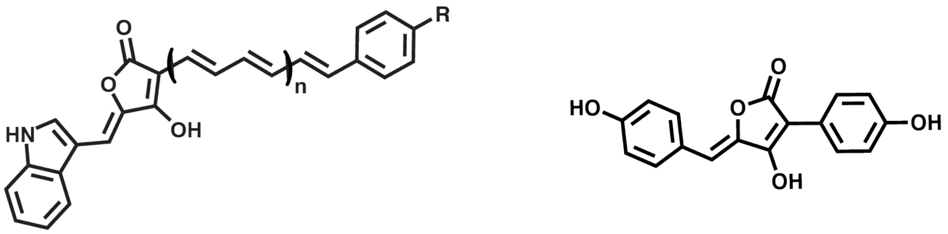

(**1**) Flavoaffinin 407: n = 1.5, R = H
(**2**) Flavoaffinin 423: n = 1.5, R = OH
(**3**) Flavoaffinin 433: n = 2, R = H
(**4**) Flavoaffinin 449: n = 2, R = OH

Aspulvinone E

**Figure S36.** Comparison of the structures of the flavoaffinins and the fungal natural product aspulvinone E.

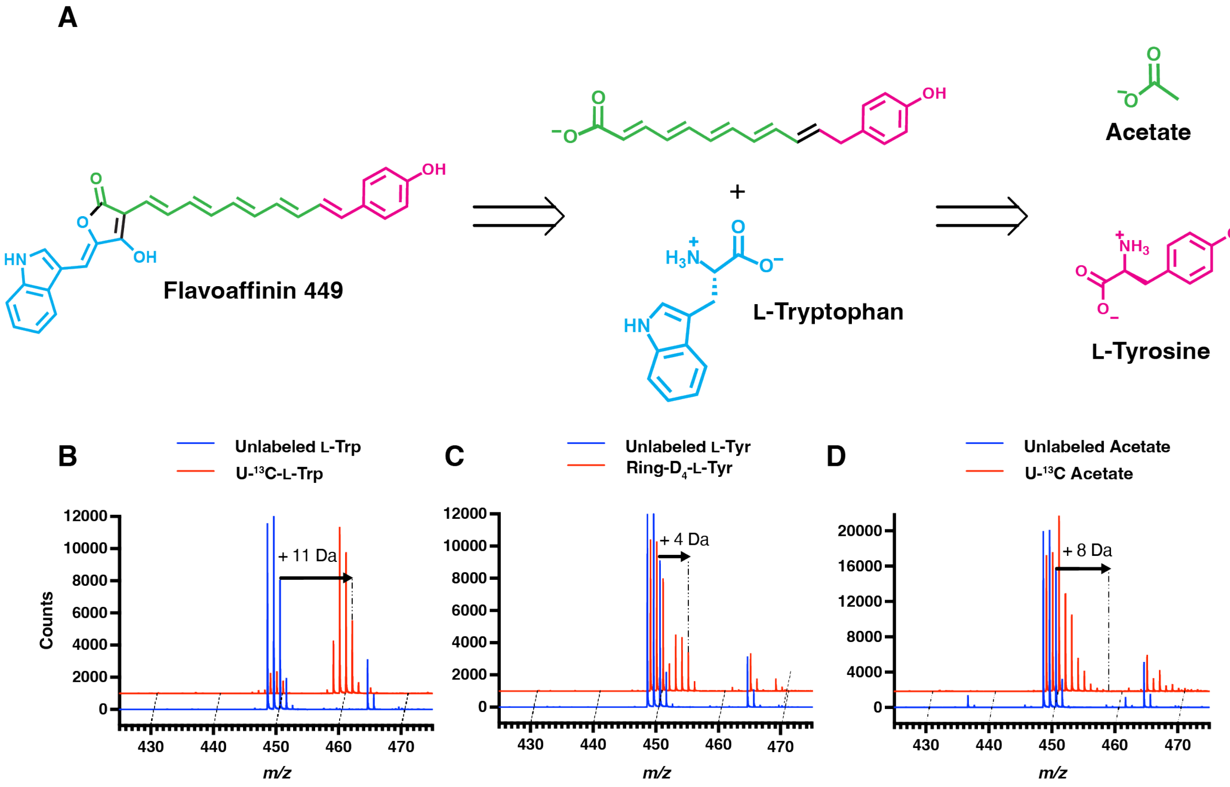

**Figure S37.** Isotopic labeling HR-ESI-MS data for flavoaffinin 449. (A) The proposed biosynthetic precursors for flavoaffinin 449. The hypothesis regarding flavoaffinin 423 is the same, but with extension by one fewer ketide. (B) HR-LC-MS spectra of *C. thermocellum* fed either 0.25 mM unlabeled L-tryptophan (L-Trp) or 0.25 mM uniformly $^{13}$C-labeled L-tryptophan (U-$^{13}$C-L-Trp). (C) HR-LC-MS spectra of *C. thermocellum* fed either 0.75 mM unlabeled L-tyrosine (L-Tyr) or 0.75 mM ring-perdeuterated L-tyrosine (Ring-$D_4$-L-Tyr). (D) HR-LC-MS spectra of *C. thermocellum* fed either 10 mM unlabeled sodium acetate (Acetate) or 10 mM uniformly $^{13}$C-labeled sodium acetate (U-$^{13}$C-Acetate). For the ESI-positive mode mass spectra in (B)-(D) each flavoaffinin analyte has ionized by multiple pathways, including as a proton adduct, as a radical cation, and following in-source dehydrogenation, as previously described in ESI-MS of other polyenes[5].

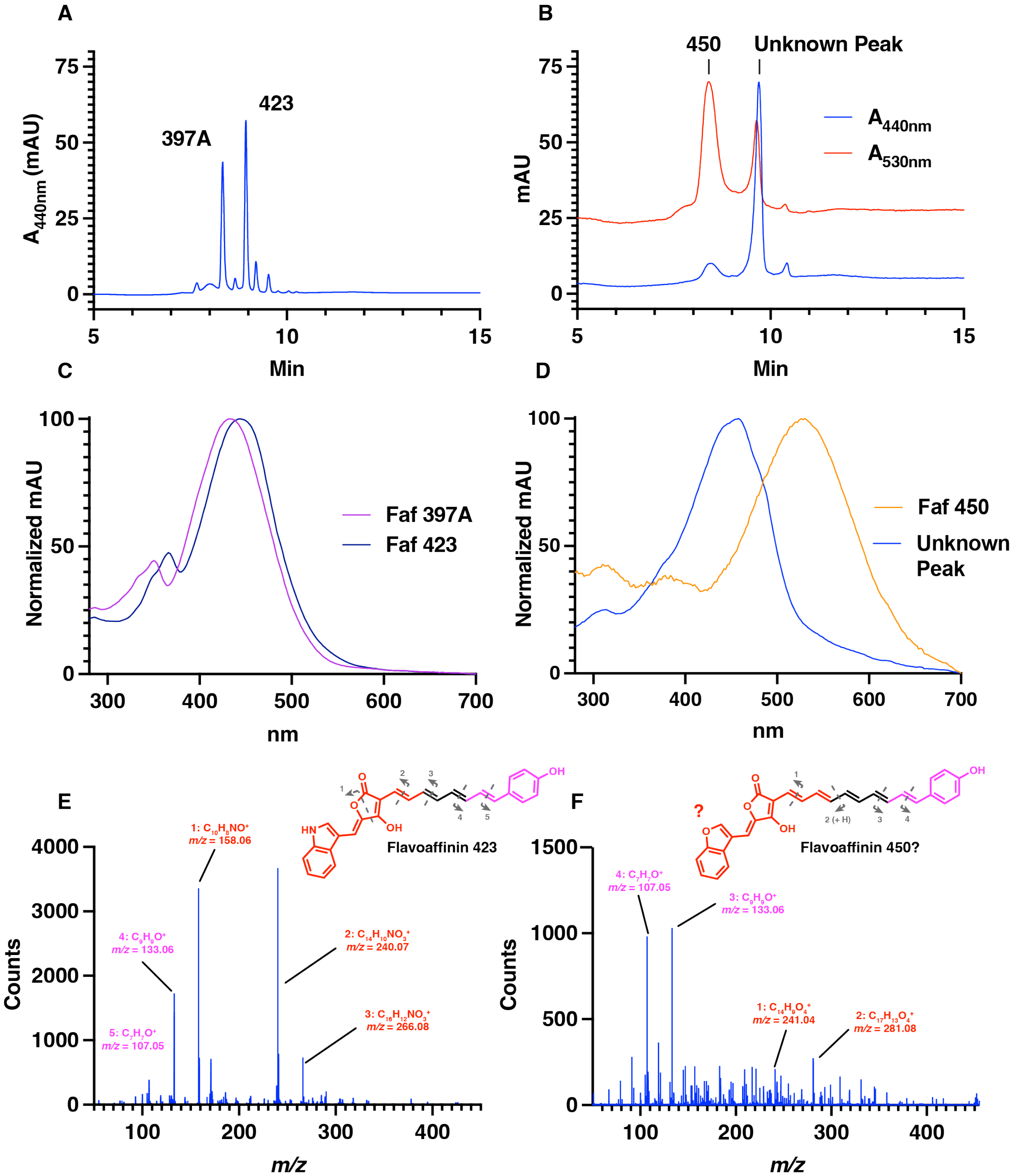

**Figure S38.** *faf* gene cluster-encoding strains of *P. cellulosolvens and R. sufflavum* produce congeners of flavoaffinins. (A) HPLC absorbance trace at 440 nm of acetone extract of *P. cellulosolvens.* Peaks corresponding to flavoaffinins 397A and 423 are labeled. (B) HPLC absorbance traces at 530 nm (red) and 440 nm (blue) of acetone extract of *R. sufflavum*. The absorbance peak corresponding to flavoaffinin 450 and an unknown prospective flavoaffinin are shown. (C) UV-Vis spectra of flavoaffinins 397 and 423 as they eluted from the HPLC column. (D) UV-Vis spectra of flavoaffinin 450 and the unknown prospective flavoaffinin as they eluted from the HPLC column. (E) MS$^2$ spectrum of flavoaffinin 423. (F) MS$^2$ spectrum of flavoaffinin 450, along with a plausible, but hypothetical structure consistent with the MS/MS data.

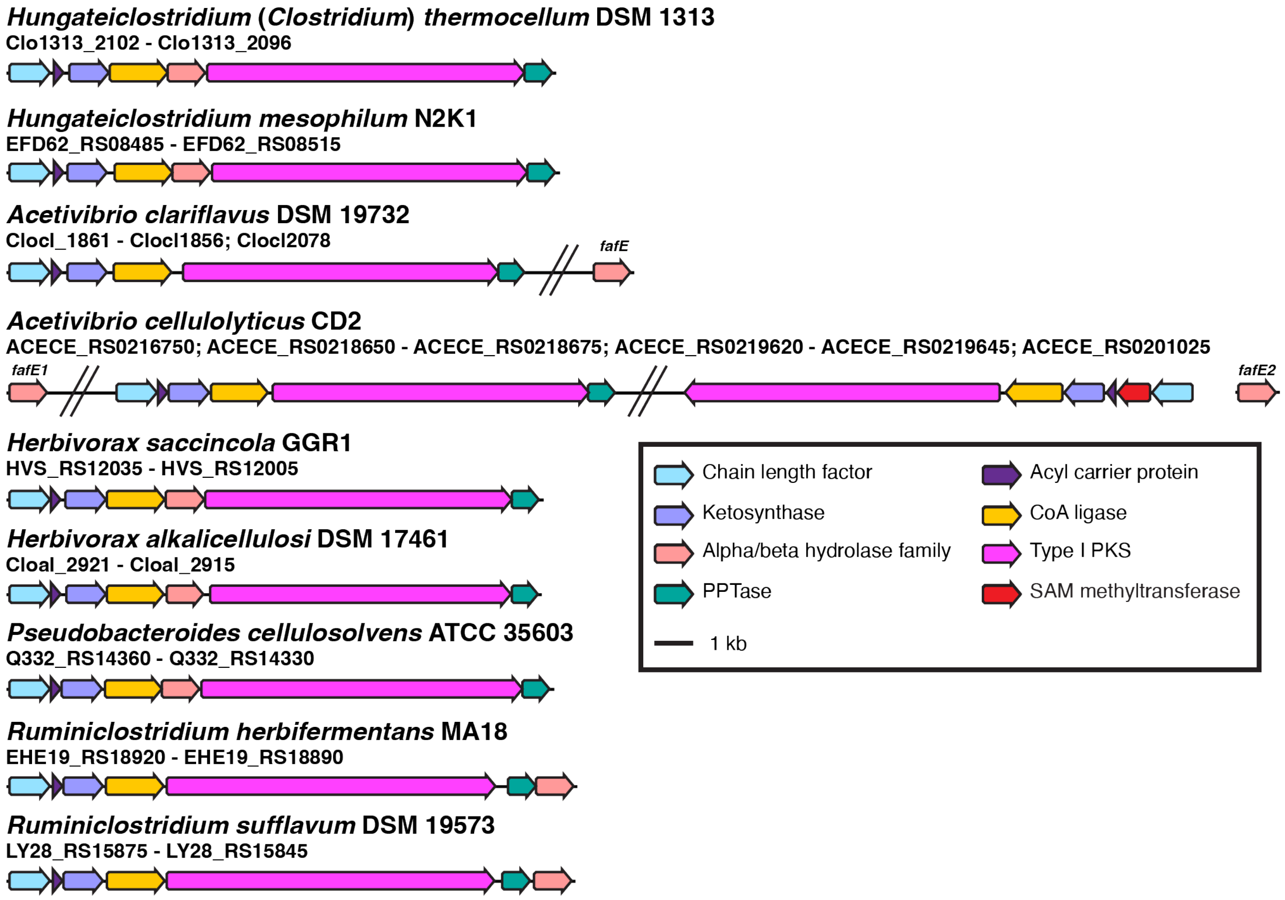

**Figure S39.** Comparison of *faf* gene clusters from other cellulolytic anaerobes. Shown are identified *faf* gene clusters, each from a species-level representative of cellulolytic anaerobe with a genome in the NCBI RefSeq database. No *faf* gene cluster-encoders were identified in this database that were not anaerobic cellulose fermenters in the order *Acetovibrionales*. Diagonal double lines indicate that a large (greater than 20 kb) DNA sequence intervenes between *faf* gene cluster components that are nonetheless present on the same assembly contig. *A. cellulolyticus fafE2* is encoded on a different contig from that harboring the other *faf* gene cluster genes and is therefore shown as a separate sequence. *A. cellulolyticus* encodes two *faf* gene clusters, one of which contains a gene annotated as encoding a SAM-dependent methyltransferase (red). The genus and species names shown are in accord with the current (as of August 2025) Genome Taxonomy Database (GTDB) taxonomy, for which *C. thermocellum* is classified as *Hungateiclostridium thermocellum*. Note that synteny within each *faf* gene cluster seems to track GTDB genus-level classification. For instance, *Acetivibrio spp.* encode FafE separately for the rest of the *faf* gene cluster, while *Ruminiclostridium spp.* encode FafE at the end, rather than middle, of the *faf* gene cluster. The relevant gene locus tags are listed below each species name. Abbreviations: PKS: polyketide synthase; PPTase: phosphopantetheinyl transferase
